# Cell type-resolved proteomics reveals intra- and intercellular signaling in Alzheimer’s disease

**DOI:** 10.64898/2026.02.02.703357

**Authors:** Xue Zhang, Kaiwen Yu, Him K. Shrestha, Ying Zhou, Jiyuan Yang, Zhen Wang, Xiaokang Ren, Ping-Chung Chen, Huan Sun, Danting Liu, Yun Jiao, Jay M. Yarbro, Ju Wang, Zhiping Wu, Kiara Harper, Liusheng He, Zuo-Fei Yuan, Xusheng Wang, Anthony A. High, Gang Yu, Jiyang Yu, Zhexing Wen, Junmin Peng

**Affiliations:** Department of Structural Biology, St. Jude Children’s Research Hospital, Memphis, TN 38105, USA; Department of Developmental Neurobiology, St. Jude Children’s Research Hospital, Memphis, TN 38105, USA; Center for Proteomics and Metabolomics, St. Jude Children’s Research Hospital, Memphis, TN 38105, USA; Department of Psychiatry and Behavioral Sciences, Emory University School of Medicine, Atlanta, GA, 30322, USA; Department of Cell Biology, Emory University School of Medicine, Atlanta, GA, 30322, USA; Department of Computational Biology, St. Jude Children’s Research Hospital, Memphis, TN 38105, USA; Core Facility of Flow Cytometry, St. Jude Children’s Research Hospital, Memphis, TN 38105, USA; BSL2 Flow Cytometry Collaborative Center, Host Microbe Interactions, St. Jude Children’s Research Hospital, Memphis, TN 38105, USA; Department of Neurology, University of Tennessee Health Science Center, Memphis, TN 38163, USA; Department of Neuroscience, Peter O’Donnell Jr. Brain Institute, University of Texas Southwestern Medical Center, Dallas, TX, USA

**Author notes:** These authors contributed equally (Xue Zhang, Kaiwen Yu, Him K. Shrestha, Ying Zhou, Jiyuan Yang).

**Keywords:** Alzheimer’s disease, neurodegenerative disease, microglia, astrocyte, oligodendrocyte, neuron, proteomics, proteome, mass spectrometry, single cell proteomics, FACS, proximity labeling, pleiotrophin

## Abstract

Alzheimer’s disease (AD) arises from pathological interactions among diverse brain cell types, but cell-specific proteomic changes remain underexplored. Here, we present deep proteomic profiling of sorted or proximity-labeled brain cells from AD mouse models (5xFAD and App^NL-G-F^) at multiple ages, quantifying 13,411 proteins in microglia (three subtypes), astrocytes, oligodendrocyte precursor cells, and neurons. We identified 3,028 differentially abundant proteins across these cell types, the majority of which were not detected in bulk proteomic datasets, and constructed cell type-specific networks to define functional modules and hub proteins. Comparison with transcriptomic data revealed that ∼30% of proteomic changes are RNA-independent. Further analyses uncovered cross-cell type signaling proteins conserved in human AD brains, such as pleiotrophin (Ptn), which is transcriptionally enriched in astrocytes but accumulates in microglia. Importantly, recombinant PTN directly activates induced microglia-like (iMG) human cells. Thus, these findings provide a comprehensive cell type-resolved proteomic atlas of AD models, highlighting novel intra- and intercellular signaling events.

**Highlights:**

- A high-resolution cell type-resolved proteomic atlas of Alzheimer’s disease mouse models
- ∼3,000 cell type-specific protein alterations identified beyond bulk tissue analyses
- Proteomic profiling of microglial subtypes reveals subtype-specific changes in Alzheimer’s disease
- Astrocyte-microglia signaling is highlighted and validated through PTN-mediated interactions

## Introduction

Alzheimer’s disease (AD) is the leading cause of dementia worldwide^1^, but the molecular mechanisms underlying its cellular heterogeneity are incompletely understood. Single-cell RNA sequencing has revolutionized AD research by uncovering disease-associated cell subtypes and cellular mechanisms^2–6^. RNA-level changes do not capture the full extent of protein-level alterations, particularly in brain tissue composed of postmitotic neurons^7–10^. Mass spectrometry (MS) has enabled extensive bulk proteome profiling of AD brain tissue, revealing large-scale protein alterations that appear to be enriched in distinct brain cell types^11–13^. However, direct cell type-specific proteomic studies in Alzheimer’s disease remain limited.

Proteomic profiling of distinct brain cell types is challenging due to difficulties in isolating specific cell types and limited protein yield from small populations^12,14^. To overcome these challenges, several strategies have been developed, including laser capture microdissection (LCM)^15^, fluorescence- or magnetic-activated cell sorting (FACS or MACS)^16^, translational labeling methods such as BONCAT^17^, and proximity labeling approaches such as TurboID^18^ or antibody-mediated biotinylation^19^ to capture cell type-specific proteomes. For instance, an early proteomic study employing MACS-MS achieved high proteome coverage of different brain cell types in mouse brain^16^. Compared with MACS, FACS provides higher purity and better resolution, particularly for microglial populations^14^. Alternatively, TurboID-based biotin labeling enabled in vivo neuronal proteome profiling through adeno-associated virus (AAV)-mediated expression^20^ or by Cre-lox transgenic mice^21^, whereas antibody-mediated biotinylation also allowed in situ cell type-specific proteome analysis in mouse brain^22^.

The mouse brain is composed of approximately 46% neurons, 20% oligodendrocytes and oligodendrocyte progenitor cells (OPC), 15% vascular cells, 12% astrocytes, 4% immune cells (mainly microglia), and 3% other cell types^23^. Despite their low abundance, microglia represent the cell population highly enriched for AD risk genes^24^. Distinct microglial subtypes are proposed by markers such as CD63 and CD74, reflecting functional reprogramming relevant to pathogenesis^25,26^. But microglial subtype-specific proteomic alterations are largely masked in current bulk tissue analyses. Advancing single cell type proteomics is critical for elucidating the molecular pathways that govern cell-specific responses in Alzheimer’s disease.

In this study, we generated a cell type-resolved proteomic atlas of two commonly used AD mouse models. Using FACS-sorted microglia (three subtypes), astrocytes, and OPC, along with AAV-based neuronal labeling, we profiled 13,411 proteins at multiple ages in 5xFAD and App^NL-G-F^ mice. We also compared our proteomics data with the results from single cell RNA-sequencing of 5xFAD brain tissue. Hundreds of differentially abundant proteins (DAPs) per cell type were identified, revealing cell type-specific pathways, RNA-protein discordance, and conserved intercellular signaling between mouse and human datasets. Pleiotrophin (Ptn) was identified to mediate astrocyte-microglia communication. This atlas highlights AD-related cellular heterogeneity and cell-cell interactions and is available at https://penglab.shinyapps.io/pannda.

## Results

### Cell type-resolved proteomics of brain cell types in AD mouse models

We developed a cell type-resolved proteomic platform that combined FACS or TurboID-based proximity labeling with 18-plex tandem-mass-tag (TMT), extensive two-dimensional liquid chromatography, and tandem mass spectrometry (LC/LC-MS/MS)^27–29^ (**Figure 1A, Figure S1**). This strategy enabled systematic profiling of four major brain cell types in two AD mouse models: APP transgenic 5xFAD (FAD) overexpressing human *APP* and *PSEN1* genes with five disease mutations^30^; and App^NL-G-F^ (NLGF) generating humanized Aβ without overexpression^31^. The analyzed cell types included three microglial subtypes (MG1-MG3 using Cd11b/Cd63/Cd74 antibodies), astrocytes (AS: Acsa2⁺), and oligodendrocyte precursor cells (OPC: O4⁺) by FACS (**Figure S2**), as well as neurons that cannot be sorted but analyzed by TurboID proximity labeling^20^. It should be mentioned that the O4 antibody recognizes sulfatide expressed in oligodendrocyte lineage cells transitioning from OPC to immature oligodendrocytes^32^; in this study, we use “OPC” to refer to this O4⁺ population.

**Figure 1.**
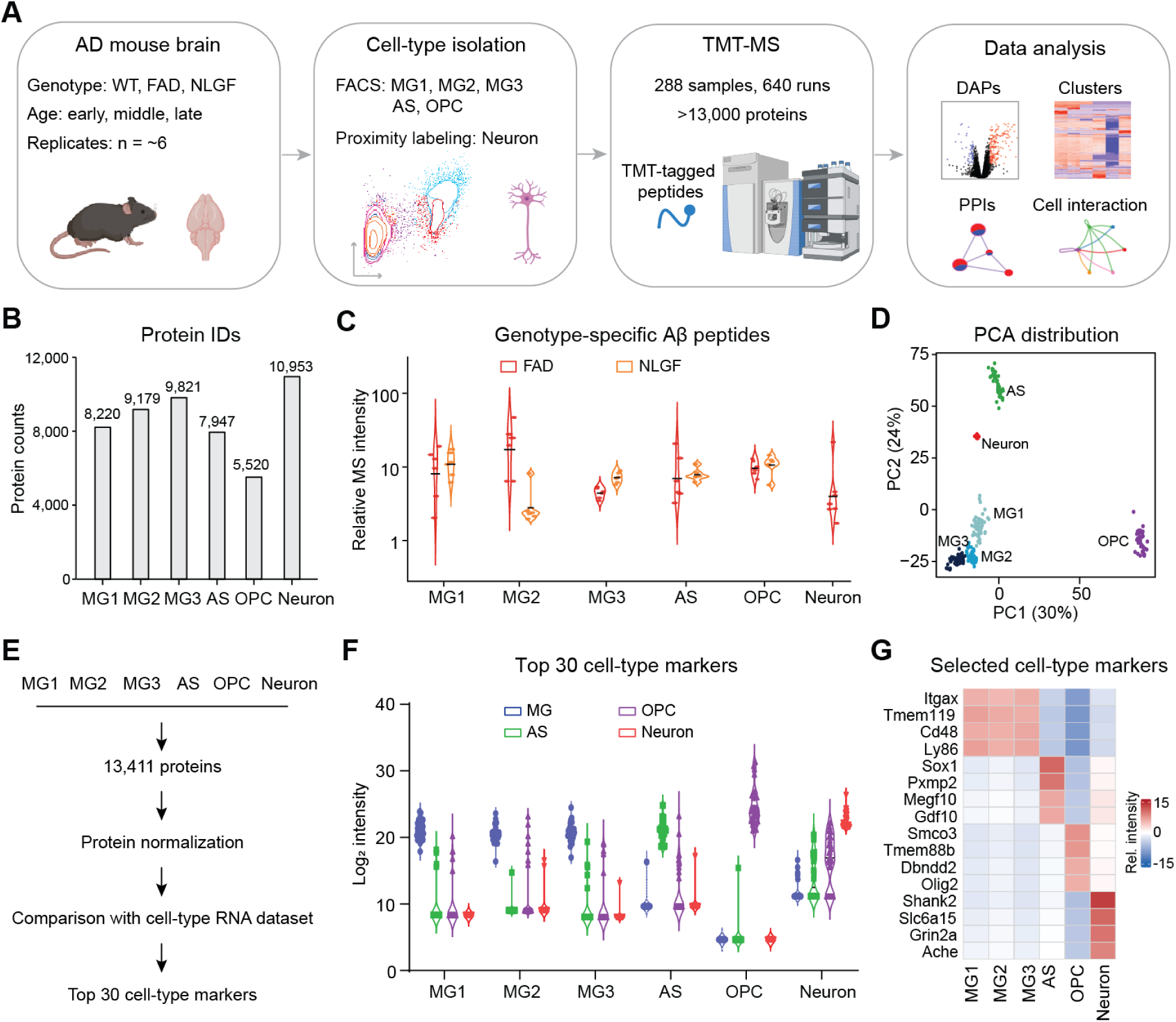
Deep *in vivo* cell type-resolved proteomic profiling of the AD mouse brains. **(A)** Schematic of the experimental workflow. Microglial subtypes (MG1, MG2, MG3), astrocytes (AS), and oligodendrocyte precursor cells (OPC) were isolated by FACS from wild-type, 5xFAD (FAD), and APP^NLGF^ (NLGF) mouse brains at early, middle, and late ages. Neuronal proteomes were profiled using proximity labeling with neuronal expression of the TurboID enzyme. **(B)** Bar plot showing the number of proteins identified across the brain cell types. **(C)** Violin plots of the relative MS intensity for genotype-specific Aβ peptides of LVFFA***E***DVGSNK and LVFFA***G***DVGSNK in FAD and NLGF mice, respectively. The MS intensities in both mice were normalized by the average signals detected in the control mice. **(D)** PCA demonstrates distinct clustering of the cell types. **(E)** Workflow for the identification of top 30 markers for each cell type. **(F)** Violin plots displaying the Log_2_ intensity of the top 30 cell type markers per cell type. **(G)** Heatmap showing the relative intensity of selected cell type-specific markers, highlighting their enrichment in specific cell populations.

We generally analyzed two mouse models and wild-type (WT) controls at different ages, with approximately six mice per age group, totaling 288 samples. The proteomic analysis was enhanced by a microgram-scale MS workflow^33^ and JUMP hybrid database searching method^34^, quantifying 8,220 proteins in MG1, 9,179 in MG2, 9,821 in MG3, 7,947 in AS, 5,520 in OPC, and 10,953 in neurons, totaling 13,411 proteins with 3,028 altered (**Figure 1B** and **Table S1,** details in the following sections). As expected, Aβ peptides were detected in these cell types from both FAD and NLGF models (**Figure 1C**). Principal component analysis (PCA) further revealed distinct proteomic profiles among cell types, with three microglia subtypes clustering closely (**Figure 1D**).

To further verify the analyzed single cell type proteomes, the results were compared to cell type specific transcriptomes^35^ used to define marker genes (**Figure 1E**). Examination of protein levels for the top 30 markers show clear and consistent cell type enrichment (**Figure 1F** and **Table S1**). For example, the microglial marker Itgax (Cd11c) were detected in three MG subtypes relative to other major cell types; the astrocytic phagocytic receptor Megf10^36^ was enriched in AS; the oligodendrocyte lineage transcription factor Olig2^37^ in OPC; and the postsynaptic scaffold protein Shank2^38^ in neurons (**Figure 1G**). Together, these results demonstrate that this cell type-resolved proteomic platform enabled deep profiling of major brain cell populations in AD mouse models.

### Proteomic alterations in three microglial subtypes

We profiled three MG subtypes at 4, 8, and 16 months of ages in the wild-type, FAD and NLGF mice (**Table S2**). In FACS, the combination of antibody selections against Cd11b (an integrin subunit encoded by Itgam, a microglial marker) ^25,26^, Cd63 (a tetraspanin associated with endosomal/lysosomal activity) ^25,26^, and Cd74 (an MHC class II subunit indicative of immune activation)^5^ enabled the separation of three microglial populations: MG1: Cd11b⁺/Cd63⁻/Cd74⁻; MG2: Cd11b⁺/Cd63⁻/Cd74⁺; MG3: Cd11b⁺/Cd63⁺ (**Figure 2A** and **Figure S2B,D**). Moreover, comparative proteomic analyses identified a subset of proteins uniquely enriched within each subtype, with the top 15 proteins shown in MG1 (such as Cyba and Fgd3), MG2 (Slc18b1 and Rbl1), and MG3 (Gpnmb and Anxa4) exhibiting strong correlations with the markers Cd11b, Cd74, and Cd63, respectively (**Figure 2B-E** and **Table S2**). Based on these subtype markers, MG1 appears to maintain a homeostatic phenotype, while MG2 and MG3 may display responsive and activated states. Consistent with microglia activation in AD, FACS analysis revealed increased numbers of all three MG subtypes in both AD mice compared with the controls (**Figure S3A**).

**Figure 2.**
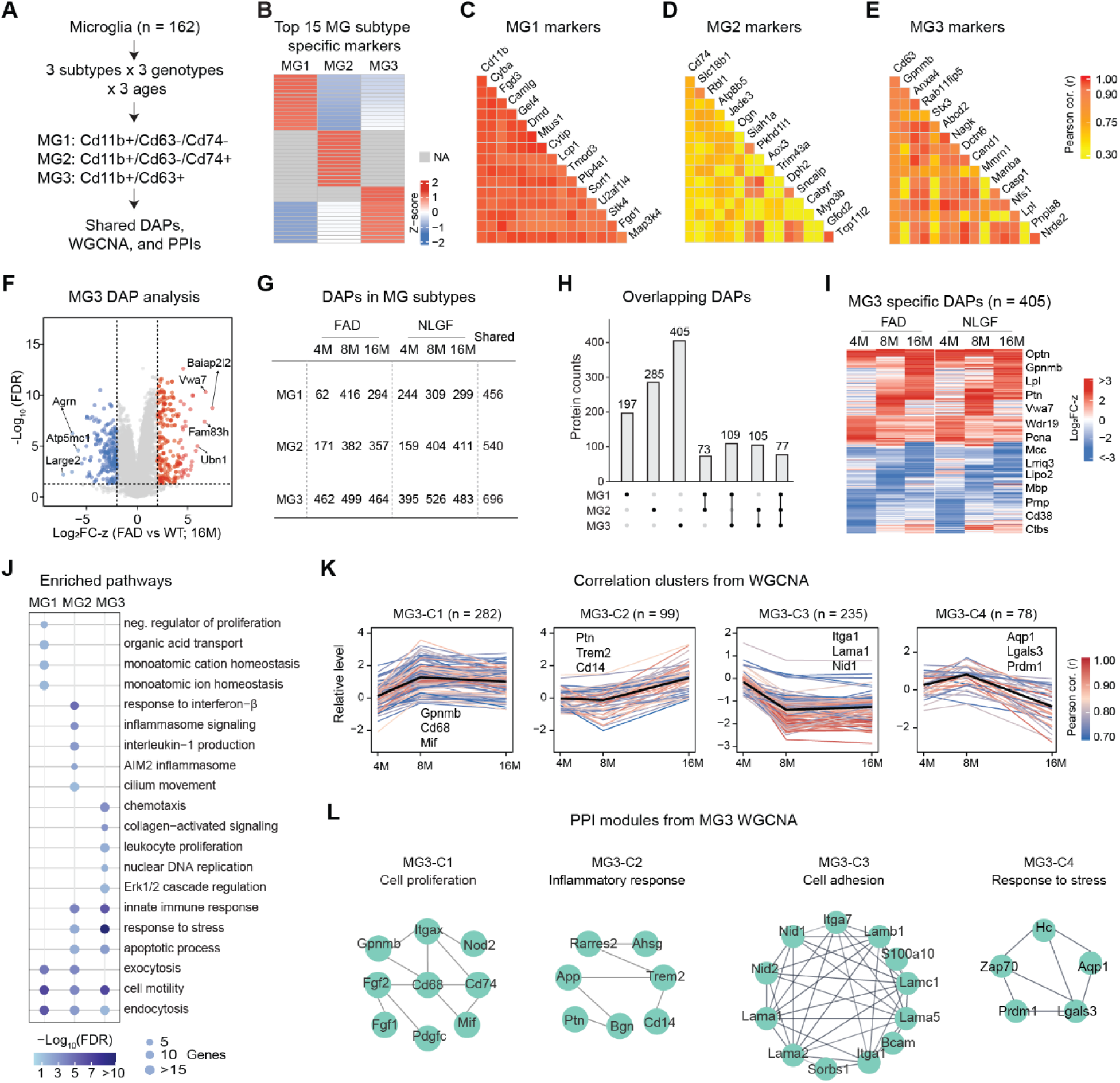
Proteomics reveals distinct molecular signatures of microglial subtypes. **(A)** Workflow for analyzing three microglial subtypes. **(B)** Heatmap of the top 15 protein markers distinguishing MG subtypes, exhibiting strong correlations with the markers CD11b, CD74, and CD63, respectively. **(C-E)** Pearson correlation heatmaps of the top subtype-specific marker proteins for MG1, MG2, and MG3. **(F)** Representative volcano plot from MG3 at 16 months comparing FAD with WT. **(G)** Numbers of DAPs identified in each microglial subtype in both AD mice. **(H)** UpSet plot showing the overlapping of DAPs among MG subtypes. **(I)** MG3-specific DAPs across ages in FAD and NLGF showing the same directional trend. **(J)** Pathway enrichment for proteins in each MG subtype; dot size corresponds to the number of genes and color intensity represents enrichment FDR. **(K)** MG3 DAP clusters identified by WGCNA; each plot shows relative protein abundance within a cluster, individual lines represent single proteins, and the black line indicates the cluster eigengene trend. **(L)** Representative PPI modules in MG3 clusters.

To examine relationships among the three MG subtypes, we performed Pearson correlation analyses using the proteome datasets across ages and genotypes. MG2-MG3 comparison showed the highest correlation, followed by MG2-MG1 and MG3-MG1, implicating MG3 as the most activated state, MG1 as homeostatic, and MG2 as intermediate (**Figure S3B**). In addition, correlations of the same subtypes between AD mice (e.g., MG3 in FAD versus MG3 in NLGF) increased with age, reflecting age-dependent microglial activation (**Figure S3C**).

To evaluate proteomic alterations within each MG subtype, we performed differentially abundant protein analysis using stringent cutoffs (AD versus WT; false discovery rate (FDR) < 0.05, |log₂FC-z| > 2; **Figure 2F** and **Table S2**) and focused on consistent changes observed in both AD models. We identified 456, 540 and 696 consistent DAPs in MG1, MG2, and MG3, respectively (**Figure 2G**). MG3 was the most responsive subtype, with 405 unique DAPs, followed by MG2 (285 DAPs) and MG1 (197 DAPs), while 77 DAPs were shared across all subtypes (**Figure 2H**). The unique DAP pattern of each subtype was largely concordant in three ages examined (**Figure 2I** and **Figure S3D,E**). The unique and shared DAPs were further analyzed for enriched pathways (**Figure 2J** and **Table S2**).

We next performed Weighted Gene Co-expression Network Analysis (WGCNA)^39^ on DAPs to delineate their temporal trajectories during disease progression at 4, 8, and 16 months. Four clusters (C1-C4) were identified within each MG subtype (**Figure 2K** and **Figure S3F**). Functional enrichment revealed distinct protein-protein interaction (PPI) modules in each cluster (**Figure 2L** and **Figure S3G**). In the case of MG3, C1 (282 proteins) exhibited an early and sustained increase, including cell proliferation proteins such as Gpnmb, Cd68, and Mif; C2 (99 proteins) showed a late increase, featuring inflammatory response proteins such as Ptn, Trem2, and Cd14; C3 (235 proteins) displayed an early decrease, including cell adhesion proteins such as Itga1, Lama1, and Nid1; and C4 (78 proteins) showed a late decrease, encompassing stress response proteins such as Aqp1, Lgals3, and Prdm1. Thus, these results indicate that amyloidosis does not induce a uniform microglial response but instead engages distinct microglial populations with specialized patterns.

In addition, we mapped all known transcription factors (TFs) and AD risk genes within the MG proteomes and DAPs. A total of 553 TFs were detected out of 1,872 curated TFs^40^, 31 of which showed consistent changes in both AD mice, including Spi1, Ikzf3, and Pparg (**Figure S3H**). Moreover, out of 167 AD risk genes^11^, 89 gene-encoded proteins were identified in MG, 24 of which were consistently altered among microglial subtypes in both mice, including Tmem106b, Clu, Trem2, and Apoe (**Figure S3I**).

Together, these analyses define three proteome-based MG subtypes that exhibit age- and disease-dependent activation, subtype-specific molecular signatures, and distinct regulatory profiles underlying their diverse responses to amyloid pathology.

### Proteomic remodeling of brain cells beyond microglia

In astrocytes, we applied a similar proteomic workflow to identify astrocyte-specific marker proteins, as well as DAPs in the FAD and NLGF models (**Figure 3A,B** and **Table S3**). Some proteins, including Adhfe1, Gcat, and Mlc1, were robustly correlated with the Acsa2 sorting marker encoded by the Atp1b2 gene^41^ (**Figure 3B**). Among 7,947 quantified proteins at the ages of 4, 8, 16 months, 613 DAPs were identified (AD versus WT; FDR < 0.05, |log₂FC-z| > 2), with 251 shared between the AD genotypes; and the “genotype-specific” DAPs also showed similar trends in both AD mice (**Figure 3C,D**). The shared DAPs were further classified into four distinct age-associated clusters (C1-C4) using WGCNA, followed by pathway and PPI module analyses (**Figure 3E-G**). C1 (59 proteins) displayed a constitutive accumulation pattern associated with cellular stress adaptation, including Kcnj8 and Hspb1 (Hsp27). C2 (20 proteins) reflected a late cytoskeletal response, characterized by actin-sequestering proteins such as Tmsb4x, Tmsb10 and Tmsb15a. C3 (123 proteins) exhibited an early decline from 4 to 8 months, featuring synaptic components of Snap25, Sv2a and Bsn. C4 (49 proteins) had a late-stage attenuation pattern enriched for interferon alpha response factors (Ifit3b, Ifit3, and Ifit1), suggesting a shift to immune activation as AD progresses. We also analyzed TFs and AD risk genes within the astrocytes, 10 of which were consistently altered in both AD mouse models, including Tbx2, Tbx3, Mnda, and Mapt (tau) proteins (**Table S3**).

**Figure 3.**
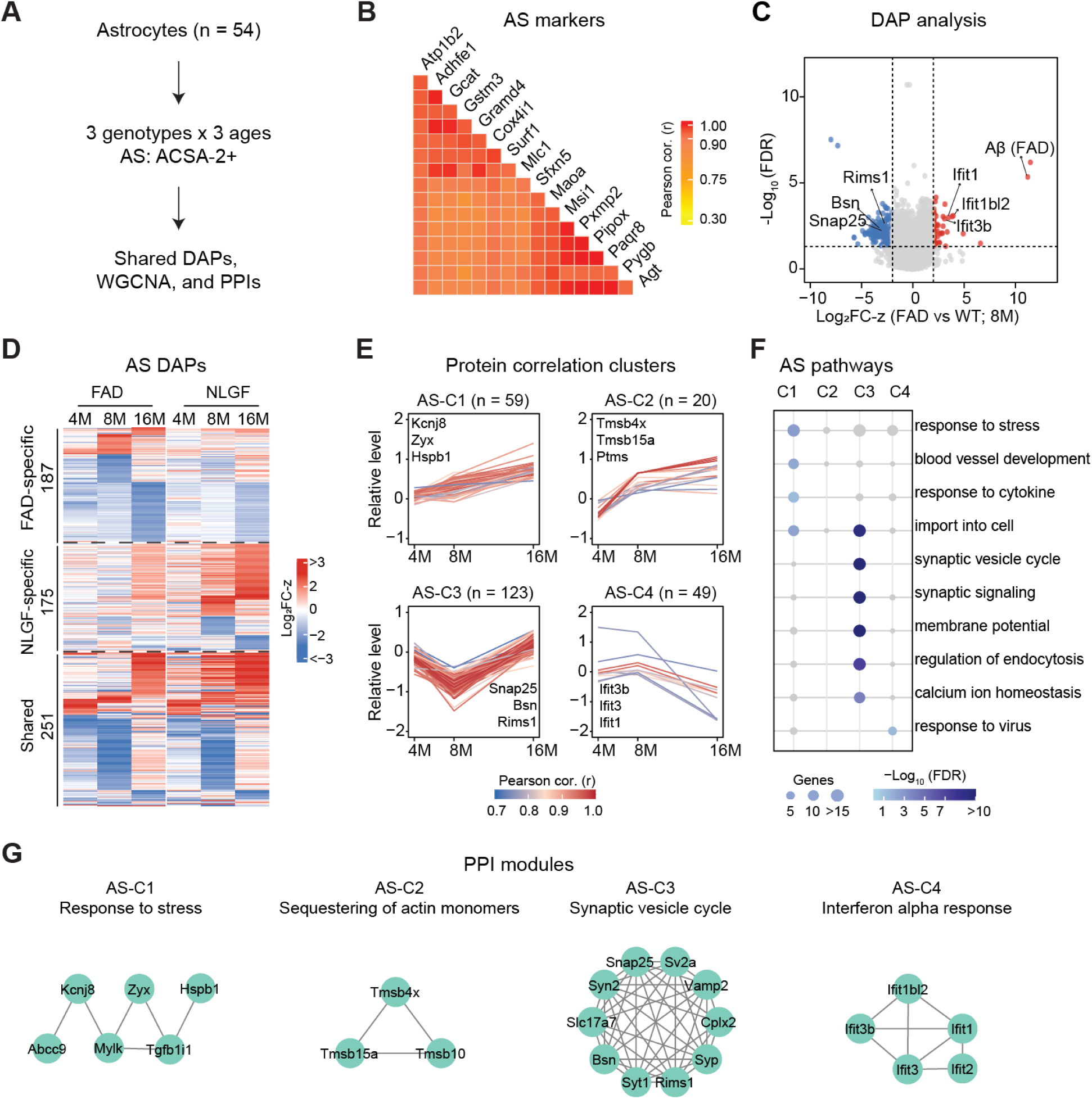
Proteomic characterization of astrocytes in AD models. **(A)** Workflow of astrocyte analysis. **(B)** Pearson correlation heatmap of top AS-specific marker proteins, ranked by their correlation with Atp1b2, the target of the ACSA-2⁺ antibody. **(C)** Representative volcano plot from astrocyte at 8 months comparing FAD with WT. **(D)** Heatmaps of age-dependent DAPs grouped by genotypes: FAD-, APPKI-specific, and shared by both models. **(E)** Four astrocyte protein clusters identified through WGCNA. Each line represents a single protein’s trajectory over time. **(F)** Pathway enrichment of the four astrocyte protein clusters. Dot size corresponds to the number of overlapping genes, and color intensity reflects the statistical significance. **(G)** PPI networks of enriched biological processes from WGCNA modules.

In oligodendrocyte precursor cells, we identified OPC-specific marker proteins and DAPs in the FAD and NLGF models (**Figure S4A,B** and **Table S4**). OPC markers such as Ugt8, Prr18, and Gjb1 were detected (**Figure S4B**). A total of 5,520 proteins were quantified, in which 371 DAPs were found (AD versus WT; FDR < 0.05, |log₂FC-z| > 2), with 116 shared between the AD genotypes (**Figure S4C,D**). The shared DAPs were further classified into two distinct clusters (C1-C2) and subjected to pathway and PPI analyses (**Figure S4E-G** and **Table S4**). C1 (46 proteins) showed a constitutive accumulation pattern associated with cell differentiation and neuroinflammatory responses, including Gap43, Naglu, and Snca. C2 (69 proteins) reflected a down-regulation related to RNA splicing and nucleosome assembly. RNA splicing defects have been consistently shown to accelerate cognitive decline in the presence of β-amyloid aggregation^42,43^. In addition, seven TFs showed consistent changes in both models, including Hmga1, Stat6, and Hif3a (**Table S4**). The proteins encoded by four AD risk genes, including App, Agrn, Grn, and Tmem106b, were consistently altered in the OPC (**Table S4**).

Given the limited efficiency of neuronal isolation by FACS, we analyzed control and FAD mice at 12 months of age using AAV-mediated, CaMKIIα promoter-driven expression of TurboID for neuronal proteome labeling^20^. Among 10,950 detected proteins, 9,462 were enriched (AAV versus non-AAV; FDR < 0.05, log₂FC > 0), and 97 showed significant changes (AAV-AD versus AAV-WT; FDR < 0.05, |log₂FC-z| > 2, **Figure S5A-C** and **Table S5**). Pathway analysis revealed that upregulated DAPs were associated with lipid metabolic processes (e.g., Srebf2, Lpcat3, and Cyp7b1) and homeostatic process (e.g., Lamp, Tmem106b, and Tfrc), whereas downregulated DAPs were primarily enriched in neurotransmitter transport (e.g., Slc17a6, Sncg, and Chat) (**Figure S5D,E**).

In summary, these findings indicate coordinated and cell-type-specific proteomic remodeling in astrocytes, OPC, and neurons, revealing dynamic stress responses, inflammatory activation, synaptic decline, and alterations in key transcription factors and AD risk genes/proteins in these AD mouse models.

### Cell type specific modules and hubs revealed by network Analysis

To further explore cell type-specific molecular drivers, we used all proteins identified across cell types to construct a co-expression network using the MEGENA program^44^ (**Figure 4A** and **Table S6**). MEGENA revealed that the resulting modules contained proteins derived from multiple cell types, highlighting both shared and cell type-specific contributions (**Figure 4B,C**). We then mapped DAPs onto these networks to identify affected modules within the global co-expression landscape (**Figure 4D** and **Table S6**).

**Figure 4.**
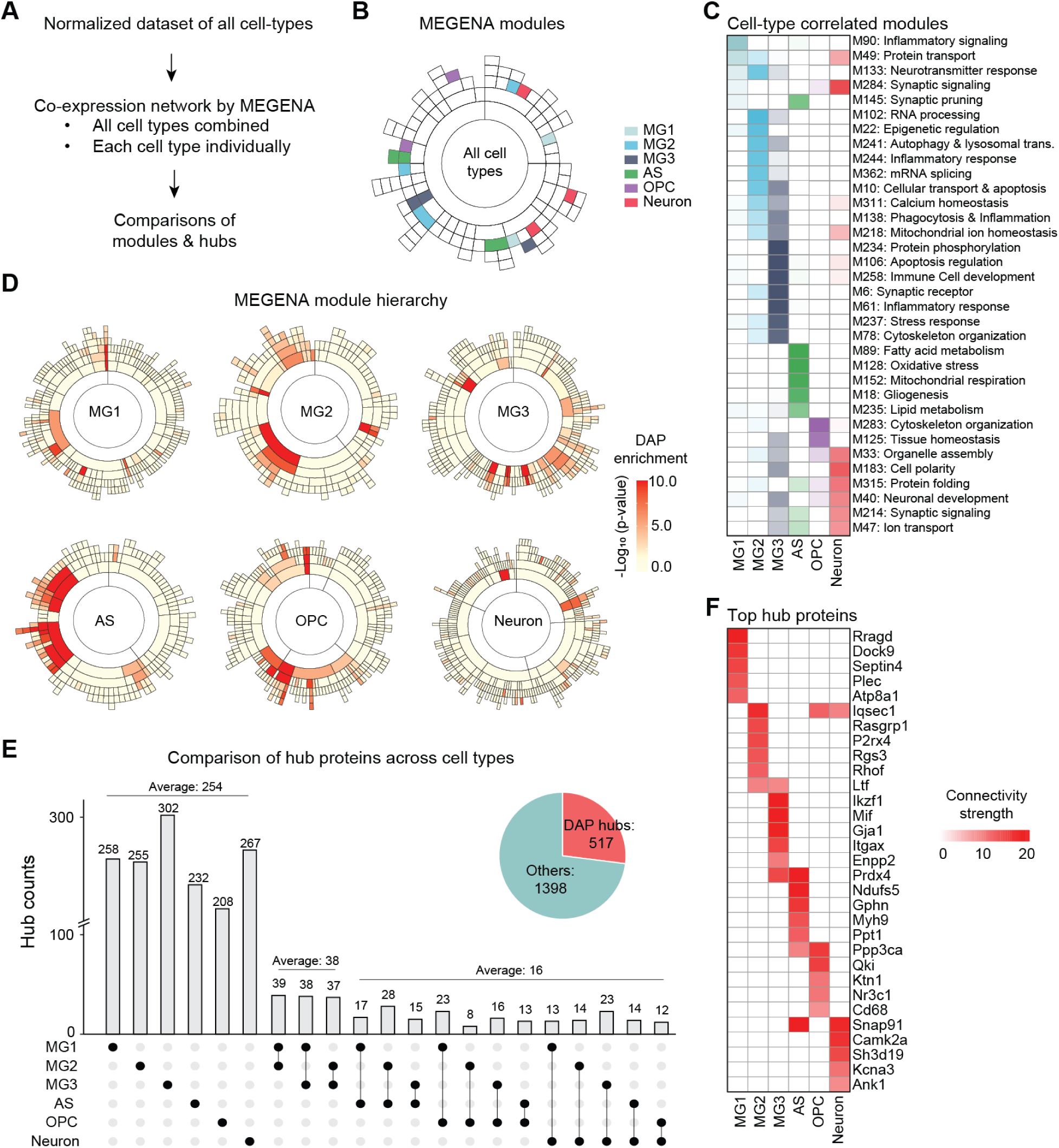
Co-expression network analysis identifies functional modules from cell types. **(A)** Workflow for module and hub identification using MEGENA using all identified proteins. **(B)** Sunburst plot showing modules identified, colored by their association with MG1, MG2, MG3, AS, OPC, or neurons. **(C)** Heatmap of module-cell type relationships, where color intensity represents the correlation between each module and cell type. **(D)** Module hierarchies for individual cell types, with color intensity indicating enrichment of DAPs within each module. **(E)** UpSet plot comparing hub proteins from modules in cell types. The inset pie chart indicates the proportion of DAPs among the identified hub proteins. **(F)** Heatmap of the module node strength of the top hub proteins across cell types.

Next, we analyzed hub proteins in each cell type, where a hub is defined as a protein with high topological connectivity (e.g., node degree or centrality) within a co-expression module of the multiscale network, potentially representing key drivers critical for network organization and cellular phenotypes. In total, 1,915 hubs were identified in the global MEGENA network, of which 517 overlapped with DAPs (**Figure 4E**). Each cell type contained at least 200 unique hub proteins.

As expected, microglial subtypes shared an average of 38 hub proteins with one another, substantially higher than the average of 16 hubs shared among other cell types.

Among these hub proteins, top highly connected proteins emerged as central regulators of pathogenesis in a cell type-specific manner (**Figure 4F**). For example, in MG1, Atp8a1, a P4-ATPase lipid flippase essential for maintaining phospholipid asymmetry and vesicle/membrane homeostasis^45^, emerged as a central hub. In MG2, P2rx4 (P2X4 purinergic receptor), which promotes ApoE degradation^46^, was strongly enriched, whereas MG3 was dominated by Mif (macrophage migration inhibitory factor)^47^, a pro-inflammatory cytokine that drives microglial activation. In astrocytes, the key lysosomal depalmitoylating enzyme Ppt1 was identified. Ppt1 directly depalmitoylates GFAP and serves as an important regulator of astrocyte reactivity and gliosis^48^. OPC was enriched for RNA splicing hubs, with Qki, a master regulator of oligodendrocyte differentiation and myelin gene expression^49^, standing out as a key driver of white-matter integrity. In neurons, hubs such as Snap91 and Camk2a, key regulators of synaptic vesicle cycling and synaptic plasticity^50,51^, reflecting the vulnerability of synaptic pathways in AD.

Overall, MEGENA network analysis identified cell type specific modules, hub proteins central to these networks, and DAP-associated hubs, revealing distinct but interconnected molecular programs potentially contributing to AD pathogenesis.

### Protein Changes Masked in Bulk but uncovered by Single Cell Type Analysis

We compared this dataset with mouse brain bulk proteomic results^10^ to determine which DAPs were hidden in bulk studies but discovered by single cell type proteomics (**Figure 5A** and **Table S7)**. After summing all cell type proteomes, 9,027 proteins were shared between the bulk and cell type datasets (>12-month-old mice, **Figure 5B** and **Table S7**). At the individual cell type level, ∼25% of proteins identified by cell type proteomics were absent from bulk analyses **(Figure 5C**). For cell type DAPs, the proportions missed in bulk analysis were 35% in neurons, 61% in AS, 77% in OPC, and 83-86% in MG (**Figure 5D**). Overall, the majority (78%) of cell type DAPs were not captured in bulk (**Figure 5E** and **Table S7**). Thus, cell type-resolved analysis provides a high-resolution view of the proteomic landscape.

**Figure 5.**
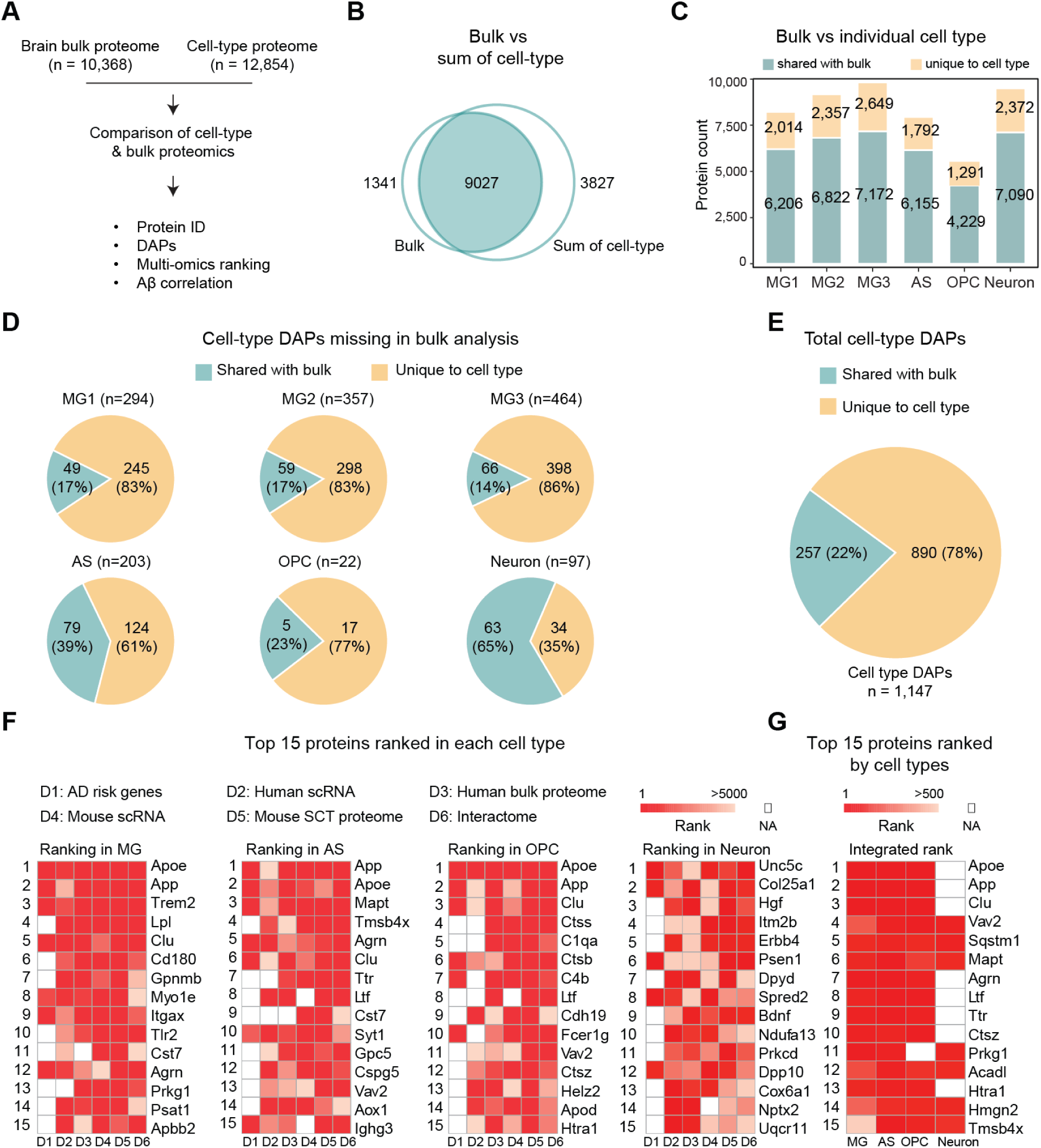
Comparison of cell type and bulk brain proteomics in AD mice and multi-omics ranking. **(A)** Workflow for comparing cell-type and bulk proteome in FAD mice. **(B)** Overlap of proteins detected by the two methods. **(C)** Protein counts per cell type, highlighting proteins shared with bulk versus unique to cell type proteomics. **(D)** Proportion of cell-type DAPs (≥12 months) present or absent in bulk (blue: shared; yellow: missing). **(E)** Summary of 1,147 cell type DAPs (≥12 months): 22% shared with bulk and 78% only in single cell type proteomics. **(F)** Multi-omics ranking using order statistics across six datasets: (1) AD risk genes; (2) human scRNA-seq; (3) human bulk proteome; (4) mouse snRNA-seq; (5) mouse cell-type proteome; and (6) AD core interactome. Heatmaps show the top 15 ranked proteins per cell type. **(G)** Integrated ranking of the top 15 proteins based on summed ranks across all cell types.

To prioritize DAPs relevant to disease development, we integrated multi-omics data to rank DAPs in each cell type across six datasets, including mouse cell type proteome (this study), mouse single cell transcriptome^35^, GWAS^11,12^, human bulk proteome (no human cell type proteomic data available), human single cell transcriptome^3,4^, and protein-protein interactome^8^ (**Figure 5F** and **Table S7**). In microglia, the highest ranked proteins included Apoe, Trem2 and Lpl, reflecting lipid sensing and clearance. In astrocytes, Mapt, Tmsb4x and Agrn ranked at the top, capturing cytoskeletal dynamics, actin regulation and extracellular matrix organization. In OPC, C1qa, Ctss and C4b were most enriched, indicating complement activation and antigen processing functions. In neurons, Unc5c, Col25a1 and Psen1 ranked highest, reflecting the Aβ processing pathway.

Furthermore, the overall ranking based on all cell types above (**Figure 5G**) across genetic, transcriptomic and proteomic datasets consistently positioned Apoe, App, and Clu among the top candidates, together with emerging factors such as Agrn, Ltf and Ttr. In addition, we analyzed the DAPs correlated with the levels of Aβ, identifying the top responsive DAPs of Ifit1, Cd68 and Apoe (**Figure S6** and **Table S7**).

### Intercellular signaling proteins revealed by RNA-protein inconsistency

The inconsistency between transcriptomic and proteomic changes in human AD brains^8,9^ and mouse models^10^ have been reported previously in bulk analyses. To extend this comparison to individual cell types, we compared the cell-type proteomes with our cell-type transcriptomes^35^ in the FAD mice (**Figure 6A** and **Table S8**). Using age-matched data from ∼12-month-old mice maintained under the same breeding conditions, we found that some protein changes (32% on average, **Figure 6B**) were independent of RNA levels using the published criteria^10^. The proportions of RNA-independent protein changes were 39% in MG (combining all three subtypes), 17% in AS, 34% in OPC, and 38% in neurons. These results indicate the complementarity of transcriptomic and proteomic datasets in profiling different cell types of disease tissues.

**Figure 6.**
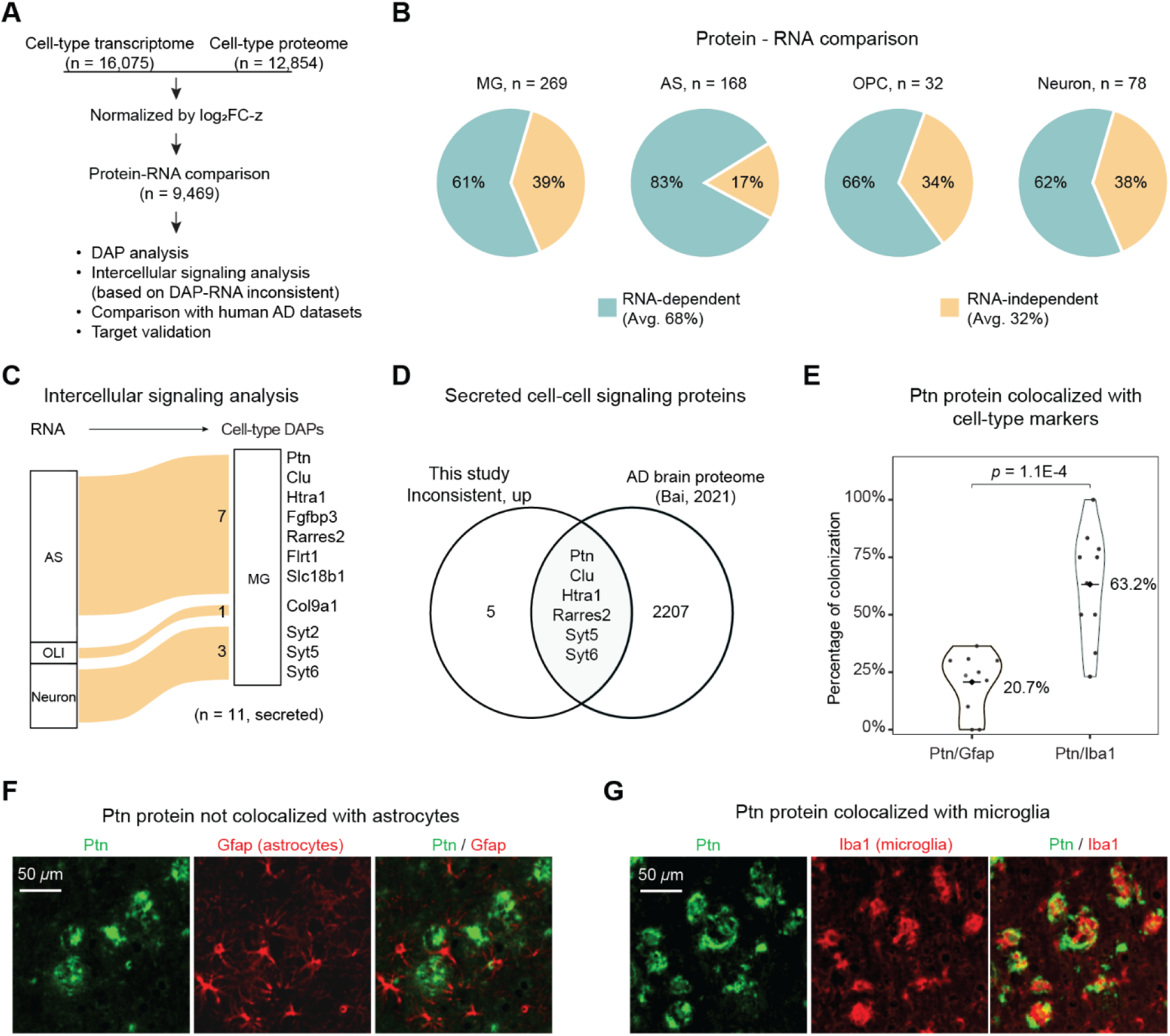
Integrative proteo-transcriptomic analysis identifies RNA-independent proteomic changes. **(A)** Workflow for comparing cell type proteome with single cell transcriptome in FAD mice (∼12 month-old). **(B)** Pie charts comparing the Protein-RNA DAPs to determine the RNA-dependent and independent protein changes. **(C)** River plot showing intercellular mismatches between protein and mRNA localization, focusing on secreted proteins. **(D)** Venn diagram identifying key proteins with potential involvement in cell-cell signaling. For this study set, proteins were filtered to contain only i) DAPs, ii) RNA-independent iii) upregulated, and iv) secreted proteins. **(E)** Violin plots quantifying the percentage of PTN colocalized with Gfap^+^ astrocytes or Iba1^+^ microglia. The Y-axis shows percentage of colocalization. **(F,G)** Immunofluorescence images of FAD mouse cortex showing Ptn (green) protein not colocalized with the astrocyte marker Gfap (red,F) but colocalized with microglial marker Iba1 (red,G). Scale bar, 50 µm.

RNA-protein inconsistency can arise from several factors: (i) intracellularly, from post-transcriptional regulatory mechanism such as altered protein degradation rates^10^; (ii) intercellularly, when proteins produced in one cell type are secreted and subsequently taken up or bound by another cell type, leading to mismatches between RNA and protein levels within individual cell types. We searched for molecules in the second category that may act as intercellular signaling regulators. Interestingly, 11 microglial DAPs corresponded to RNA transcripts that were enriched in astrocytes, oligodendrocytes, and neurons, suggesting their potential roles in intercellular communication between microglia and other cell types in the brain (**Figure 6C**).

To determine whether these 11 putative intercellular signaling proteins are relevant to human AD pathology, we compared the mouse data with a compiled human AD brain proteomic dataset^11^. This analysis confirmed six proteins consistently upregulated in human AD brains, including Ptn, Clu, Htra1, Rarres2, and Syt5 and Syt6 (**Figure 6D**). The presence of these shared protein changes across species supports that dysregulated intercellular communication and protein-level alterations are involved in AD pathogenesis.

### Astrocytic PTN-driven microglial activation

The identified Ptn is a secreted growth factor that regulates cell signaling through interactions with numerous cell-surface receptors^52^, and it is also highly correlated with Aβ levels in AD brains and mouse models in bulk studies^8^. While its transcript was predominantly enriched in astrocytes, its protein level was highly accumulated in MG3 of 5xFAD mice compared to WT (**Table S1**). However, its functional impact on microglia in the development of AD pathology is not well studied. We first validated the localizations of Ptn RNA and protein across cell types by microscopy in the AD mouse brain (5xFAD, ∼12-month-old). RNAscope and immunofluorescence analyses showed that Ptn mRNA colocalized with the astrocyte marker Gfap but not with the microglial marker Iba1 (*p* = 2.7E-7; **Figure S7A,B**), whereas Ptn protein overlapped with Iba1 rather than Gfap (*p* = 1.1E-4; **Figure 6E-G**), providing direct evidence for Ptn-mediated astrocyte-to-microglia signaling.

We next used human induced microglia (iMG) cells to examine how PTN regulates microglial signaling through proteomic profiling (**Figure 7A**). Recombinant PTN was produced and purified to homogeneity using a three-step protocol (**Figure 7B**). iMG cells were then exposed to titrated PTN concentrations (0, 30, 100, 300, or 1000 nM) followed by MS, quantifying 7,030 proteins. with 118 DAPs (FDR < 0.05, |log₂FC-z| > 2, **Figure 7C** and **Table S9**). Pathway enrichment further showed that PTN enhanced immune activation, cytokine production, and receptor recycling pathways, while reducing lipoprotein metabolism and filament organization (**Figure 7D** and **Figure S7C-E**).

**Figure 7.**
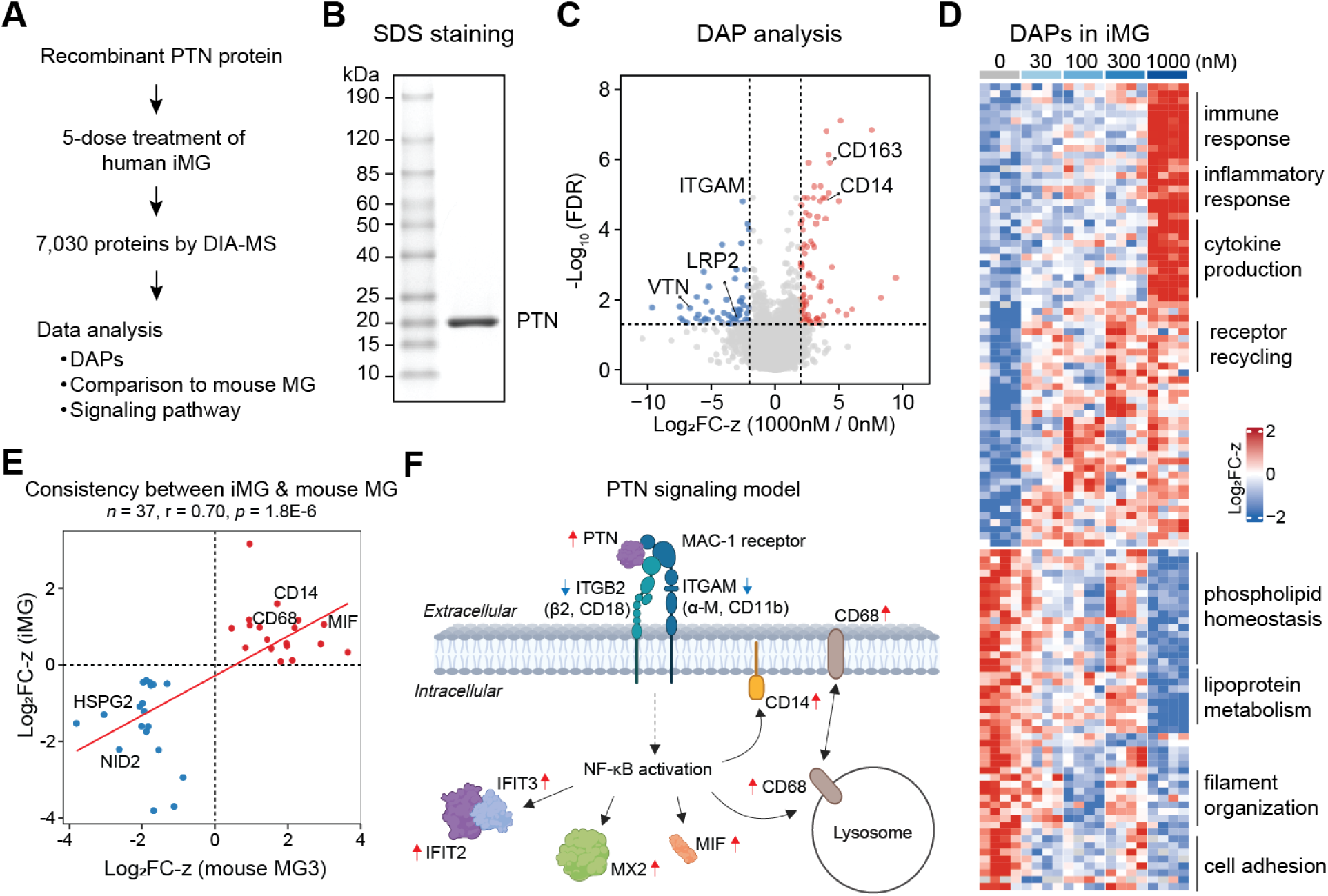
PTN stimulation alters human iMG proteome and reveals conserved signaling with mouse MG3 microglia. **(A)** Experimental workflow: recombinant PTN was purified, and applied to human induced microglia (iMG) at five doses. **(B)** Stained SDS gel confirming purified PTN. **(C)** Volcano plot of proteomic alterations induced by PTN (1000 nM vs 0 nM). **(D)** Heatmap showing dose-dependent DAP patterns across PTN treatments (0, 30, 100, 300 and 1000 nM), with functional annotations. **(E)** Scatter plot showing the correlation of PTN-induced protein changes in human iMG with mouse MG3 DAPs. **(F)** Conceptual model of PTN signaling in microglia. Arrows indicate the up-and down-regulation of proteins.

To evaluate whether these PTN-induced patterns in human iMG recapitulate microglial features in mice, we performed cross-species correlation analysis between iMG and mouse MG3 (the Ptn-enriched cell population), A positive correlation (r = 0.70) was observed with shared proteins including CD14, CD68, and MIF (**Figure 7E**). Integrating these findings and literature information^53^, we propose a PTN signaling model in which PTN may bind to the MAC-1 receptor complex (ITGAM/ITGB2)^52,54^ (**Figure S7F**), leading to the activation of the NF-κB pathway and subsequent transcriptional events. This signaling elevates expression of IFIT2, IFIT3, MX2, MIF, CD68, and CD14, driving an immune-activated microglial response (**Figure 7F**). In summary, these findings support a role for PTN as a mediator of astrocyte-microglia communication that promotes microglial activation in both mouse and human brains.

## Discussion

In this study, we generated deep AD mouse brain cell-type proteomes by profiling hundreds of samples across six cell (sub)types at three ages. In total, 13,411 proteins were quantified from microglia, astrocytes, OPC, and neurons in 5xFAD and APP-KI mice. This project was enabled by advances in highly sensitive mass spectrometry capable of analyzing minute protein amounts^33,55^, combined with classical cell-sorting strategies and the recently developed TurboID proximity-labeling approach^20,56^. These efforts establish the first comprehensive cell type-resolved proteomic atlas of AD models, revealing intra- and intercellular signaling pathways that extend beyond bulk or transcriptomic analyses.

In contrast to previous bulk studies, this work identified a large number of cell-type-specific DAPs, the majority of which were missed in bulk analyses. Notably, the proportion of missing proteins was inversely correlated with the estimated abundance of each cell type in the brain^23^ (Pearson’s r = -0.89), consistent with the idea that information from rare populations is more likely to be masked in bulk measurements. With continued advances in spatial proteomics and cell-type isolation using laser capture microdissection^10,15^, even rare cell types will become increasingly accessible for defining disease-related proteomic signatures. This analytical framework can also be extended to human samples in future studies.

RNA-protein correlation in cells is a recurring theme in multi-omics studies. Our previous comparison of bulk proteomic and transcriptomic data showed that about one-third of protein-level changes cannot be explained by RNA levels in mouse and human brains^10^. This proportion of RNA-protein inconsistency remains similar in the single-cell type analyses in this study. Intracellularly, such discrepancies may arise from alterations in posttranscriptional events, such as protein turnover. Indeed, several neuronal proteins including Abca3, Tmem63c, and Arfgef3, exhibited significantly prolonged half-lives in AD mouse brains^10^. These stabilized proteins may contribute to disruptions in essential cellular processes, including lipid homeostasis (Acba3)^57^, ion balance (Tmem63c)^58^, and vesicular trafficking (Arfgef3)^59^, implicating impaired proteostasis and pathway dysfunction.

Remarkably, microglia displayed a higher proportion of DAPs independent of RNA levels than other cell types, accounting for nearly half of all RNA-independent proteins. This may reflect their phagocytic role in engulfing protein-rich synaptic and debris from other cell types. For example, elevated presynaptic vesicle-associated proteins, such as Syt2, Syt5, and Syt6, and the autophagy regulator Atg4d, support the idea that microglia internalize presynaptic boutons through mechanisms such as trogocytosis^60^.

More intriguingly, the RNA-protein comparisons provide direct proteomic evidence for cell-cell communication, revealed by transcripts upregulated in one cell type but corresponding proteins accumulating in another. For instance, several astrocyte-secreted proteins, including Ptn, Rarres2, Clu, and Htra1, accumulated at microglia, indicating active uptake. Prior studies suggest that PTN treatment alters microgliosis and NF-κB signaling in cultured BV2 microglial cells^61^. Consistently, our proteomic profiling of PTN-treated iMG supports a model in which external PTN contributes to microglial innate immune activation, as reflected by increased Cd14, Cd68, Mif, Mx2, and interferon-related proteins, reminiscent of mouse MG3 activation. Ptn may signal through Mac-1 (Itgam/Itgb2) and Ptprz1 to regulate immune activation^53,54^. In addition, Rarres2 (chemerin) is a secreted adipokine^62^ that regulates metabolism, inflammation, and tissue homeostasis through its receptor Cmklr1 that was also detected in MG3. These findings highlight a communication axis from astrocytes to microglia, mediated by secreted proteins, which shapes microglial activation during AD progression.

Finally, we mapped known TFs and AD risk genes to our DAP data from different cell types. Coordinated transcriptional reprogramming can distinguish microglial subtype-specific functions. A recent study reveals that PU.1 (Spi1) downregulation, previously considered a hallmark of microglial dysfunction, actually drives a neuroprotective, lymphoid-like microglial phenotype that compacts amyloid plaques and reduces neuronal damage^63^. Broad downregulation of Spi1 protein in MG2 also suggests that MG2 adopts an anti-inflammatory, aggregate-processing state. Our cell type proteomics dataset provides the alterations of key TFs and AD risk genes/proteins to microglial populations, and their discrete functional consequences merit further investigation.

In summary, our study provides a deep cell type-resolved proteomic atlas of AD mouse models and reveals direct protein-level evidence of cell-to-cell communication. The dataset provides a valuable molecular framework for investigating intra- and inter-cellular mechanisms underlying Alzheimer’s disease.

## Methods

### Animals

All experiments complied with all relevant ethical regulations. Animal procedures were approved by the Institutional Animal Care and Use Committee at St. Jude Children’s Research Hospital. Three mouse models were used in this study: 1) heterozygous 5xFAD mice^30^ carrying the APP Swedish (K670N, M671L), Florida (I716V), and London (V717I) mutations and human PS1 harboring the M146L and L286V mutations; 2) homozygous APP^NL-G-F^ mice^31^ with the Swedish (K670N, M671L), Arctic (E693G), and Iberian (I716F) mutations; and 3) isogenic C57BL/6 mice. Mice at 4, 8, and 16 months of age, both female and male, were used for FACS cell isolation.

### FACS for non-neuronal cell isolation

We used previously published protocols^64,65^ with modifications. Mice were anesthetized and subjected to transcardiac perfusion with cold Dulbecco’s PBS (D-PBS without Ca²⁺ or Mg²⁺; Gibco) to remove blood. The brain was dissected on ice, transferred into a tube containing 5 mL of cold Hank’s Balanced Salt Solution (HBSS, without Ca²⁺ or Mg²⁺; Gibco) with 0.5% BSA, and excised into ∼2 mm³ pieces. A brief centrifugation (300 × g for 1 min) was used to exchange the buffer to 5 mL of dissociation buffer including 50 mM PIPES, pH 7.6, 120 mM NaCl, 5 mM KCl, 0.6% (w/v) glucose, papain (10 units; Sigma-Aldrich), DNase I (500x; 25 mg/ml stock; Roche), and 1× penicillin/streptomycin (Gibco). Brain tissues were further dissociated using a gentleMACS™ Octo Dissociator (Miltenyi Biotec) and incubated at 37 °C for 20 min. The digested tissue was passed through a 70-µm cell strainer, washed twice with cold HBSS, and washed once with D-PBS. The resulting pellet was resuspended in 5 mL of 22% Percoll (Cytiva) and centrifuged at 500 × g for 20 min with low brake. Cells at the bottom were collected, washed with HBSS containing 0.5% BSA, and resuspended in FACS sorting buffer (D-PBS with 0.5% BSA) for staining.

Brain cells were stained with a 1:200 dilution of Cd11b-APC-Cy7 (BioLegend), Cd74-Alexa Fluor 488 (BioLegend), Cd63-PE/Cy7 (BioLegend), Acsa-2-PE (Miltenyi Biotec), O4-APC (Miltenyi Biotec), and DAPI (1 µg/mL) in the dark for 20 min. The cells were then centrifuged down at 800 × g for 3 min, washed once with D-PBS, and resuspended in 500 µL of FACS sorting buffer. The suspension was passed through a 50-µm strainer and sorted on a FACS Aria III SORP using BD FACSDiva™ software. Single, viable cells were first gated, and five cell populations were isolated based on cell type-specific markers. Sorted cells were centrifugated at 800 × g for 3 min and stored at -80°C for further analysis.

### AAV-mediated proximity labeling of neuronal proteins

Recombinant adeno-associated viruses (AAVs) were generated following previously published protocols^20^. Briefly, the AAV expressing 3xFLAG-TurboID-mCherry under the CaMKII promoter were injected into 5xFAD mice and littermate controls (∼12-month-old) via the retro-orbital sinus, while a non-AAV control group was injected with saline. After three weeks, mice received one week of daily subcutaneous biotin injections, followed by tissue harvest. The cortex tissue was lysed, and biotinylated proteins were purified with streptavidin beads, and digested for TMT labeling following established protocols^20^.

### TMT-based cell type proteomics by LC/LC-MS/MS

Cells were lysed, digested, and TMT-labeled following a previously published microgram-scale proteomics protocol^33^, with minor modifications. Briefly, sorted cells were lysed in lysis buffer (50 mM HEPES, pH 8.5, 8 M urea, 0.5% sodium deoxycholate, and 1 mM DTT), with a protein yield of ∼1-10 µg per cell type from one mouse brain. Proteins were digested by LysC and trypsin. Peptides were subsequently reduced by DTT and alkylated using iodoacetamide, desalted, and dried by SpeedVac. The dried peptides were labeled with TMTpro reagents^29^, equally pooled, fractionated by basic pH LC to collect 40 concatenated fractions.

The peptides in each fraction were analyzed with acidic pH LC (C18, 1.7 µm, 75 µm × 20 cm), coupled with a Q Exactive HF Orbitrap MS (Thermo Fisher Scientific) in data-dependent mode of cycling one MS1 scan and 20 MS2 scans. The MS1 parameters were set as follows: 460∼1600 m/z, 60,000 resolution, 1×10^6^ automatic gain control (AGC), and a maximal ion time of 50 ms. For MS2 scans, the parameters were: fixed first mass of 120 m/z, 60,000 resolution, 1×10^5^ AGC, maximum ion time of ∼110 ms, isolation window of 1.0 m/z with 0.2 m/z offset, 10 s dynamic exclusion, and higher-energy collision dissociation (HCD) with a normalized collision energy (NCE) of 30%.

### Protein identification and quantification with JUMP software

Protein identification and quantification were performed using the JUMP search engine as previously described^8,20,34^. Spectra were searched against a mouse protein database combining Swiss-Prot, TrEMBL, and UCSC entries (59,423 proteins)^20^ concatenated with a reversed decoy database^66^. Search parameters included dynamic methionine oxidation (+15.99492 Da), static TMT labeling on peptide N-termini and lysine residues (+304.20715 Da), and cysteine alkylation (+57.02146 Da). The minimum peptide length was set to 6 amino acids, up to two missed cleavages were allowed, and a maximum of three modifications per peptide was permitted. A mass accuracy tolerance of ±5 ppm was applied. Protein quantification was then performed using all identified peptides with the setting of less than 1% protein FDR.

### Differential expression analysis of identified proteins

All MS intensity data were log2-transformed for downstream analysis, and contaminant proteins were removed. Within each cell type, pre-median normalization was performed across different ages. Subsequently, WT samples across ages were used as the baseline to perform robust linear regression (RLM) using the *MASS* R package (version 7.3.65) for batch correction. Finally, all cell types were normalized to the same level by the median log_2_ intensity. For the proximity labeled neuronal proteome, the non-AAV and AAV groups were first median-normalized separately, followed by correction for endogenously biotinylated proteins^20,67^. For sorted cells, DAPs were obtained by the moderated one-way ANOVA using the *LIMMA* package (version 3.62.2) at the same age. DAPs were defined using two criteria: (i) FDR < 5%, and (ii) absolute z-scores of log_2_ fold change (FAD/WT or NLGF/WT) > 2. For the neuronal proteome, DAPs were identified in two steps: (i) enriched neuronal proteins, with log_2_ fold change (AAV vs. non-AAV) > 0 and FDR < 0.05; and (ii) DAPs within the scope of above neuronal proteins, using the same criteria above with the FDR calculated by moderated t-test in the *LIMMA* package (version 3.62.2).

### Analyses of WGCNA, pathway enrichment and PPI

WGCNA was performed using the *WGCNA* package (version 1.73)^39^. To define co-expression patterns across different ages within each cell type, only genotype shared DAPs across ages were considered. The input matrix was constructed using FAD- and NLGF-to-WT ratios. A soft-thresholding power of 16 was applied to generate a robust adjacency matrix, achieving a scale-free topology model fit (*R*² = 0.8). Correlation clusters were identified using the hybrid dynamic tree-cutting method. To reduce complexity across cell types, *minModuleSize* and *mergeCutHeight* were optimized for each cell type, resulting in consensus co-expression patterns.

For pathway analysis, selected proteins (e.g., DAPs) were used as input for GO enrichment with the *gprofiler2* package (version 0.2.3). FDR values were calculated using the g:SCS correction method implemented in the tool, and pathways were significantly enriched at FDR < 0.05. PPI networks and functional modules were detected using STRING (version 12), and then exported to Cytoscape for further analysis.

### Functional module and hub identification across cell types by MEGENA

Multiscale Embedded Gene Co-expression Network Analysis (MEGENA, version 1.3.7)^44^ was applied to identify hub proteins and functional modules across cell types. A high-hierarchy comprehensive network was constructed using all quantified proteins across cell types to detect cell type-specific modules within the global network. Subsequently, individual networks were constructed for each cell type using DAPs to identify hub genes and modules altered in AD pathogenesis. The minimum module size was set to 20 for the global network, and 10 for individual cell type networks, whereas the module p-value threshold was set to 0.05, and the hub FDR cutoff to 0.05. GO enrichment analysis of each module was performed using the *clusterProfiler* package in R, and module annotations were derived qualitatively from the top enriched GO terms.

### Bulk-cell type proteomics comparison

We compared the single cell type proteomics data with a previous dataset from FAD and NLGF mice at >12-month-old^10^. The cell type proteomics dataset was generated by combining all proteins detected in individual cell types. Next, cell type DAPs were compared with bulk to assess whether these changes were reflected at the global level. For evaluating cell type DAPs against bulk data, three criteria were applied: (i) shared changes with large effects: proteins showing the same directional change and |Z| > 2 in both bulk and cell-type datasets; (ii) shared changes with small effects: DAP proteins with an absolute ΔZ (difference in Z-scores between bulk and cell type) < 2; and (iii) all remaining cell-type DAPs were classified as not detected as DAPs in the bulk analysis.

### Prioritized ranking of AD affected genes and proteins in each cell type

To systematically prioritize key genes and proteins affected by AD in each brain cell type, we implemented an integrative ranking framework based on order statistics^8,68^. This approach consolidates multiple omics datasets into a single consensus ranking, enabling robust identification of cell type specific molecular signatures. For each cell type, multiple complementary datasets were incorporated: (i) each dataset was independently ranked based on an appropriate metric (i.e., p-values, log₂FC, or log₂FC-z) depending on data type and availability. For the interactome dataset, shortest distances between all proteins and AD-associated genes, including three causal genes (APP, PSEN1/2) and three risk genes (APOE, TREM2, and UNC5C) were computed, and an interactome ranking was derived by combining these distances. (ii) the ranked datasets were integrated using order statistics to produce a consensus rank for each gene/protein. To ensure biological relevance, the rankings were further refined to include only proteins detected in our single-cell type proteome. Finally, the prioritized gene/protein lists from all cell types were merged and re-ranked using the same approach to derive a comprehensive overall ranking.

### Analysis of protein-RNA consistency in each cell type

The comparison followed a previously published method^10^ using single-cell transcriptomics data^35^ and cell type proteomics results. To reduce complexity from subtype annotation, both proteomic and transcriptomic datasets were consolidated into primary cell types (e.g., all microglial subtypes were combined as microglia). All data were converted to Z-scores based on log2 fold change, and the altered proteins (|Z| > 2) were evaluated. If the absolute ΔZ (difference in Z-scores between protein and RNA) exceeded three, the protein-RNA pair was considered inconsistent. However, if both RNA and protein showed large changes (|Z| > 2) in the same direction, the pair was still considered consistent regardless of ΔZ.

### In situ hybridization and immunofluorescence

Formalin-fixed paraffin-embedded brain sections (10 µm) were processed using the RNAscope Multiplex Fluorescent Reagent Kit v2 (Bio-Techne). Briefly, sections were deparaffinized, rinsed with isopropanol, subjected to target retrieval, treated with hydrogen peroxide to quench endogenous peroxidase activity, and partially digested with Protease Plus. RNAscope probes targeting Ptn mRNA or control probes (negative control and positive control) were hybridized on sections for 2 h at 40°C, followed by sequential amplification steps (AMP1-3) and HRP-based detection. Signals were developed using TSA Plus fluorophores (Opal 650, Akoya Biosciences) for 30 min at 40°C. For subsequent immunofluorescence, sections were blocked in 5% normal donkey serum and incubated overnight at 4°C with the following primary antibodies: anti-PTN (1:50, Santa Cruz Biotechnology), anti-IBA1 (1:100, FUJIFILM Wako), and anti-GFAP (1:100, Synaptic Systems). Corresponding Alexa Fluor-conjugated secondary antibodies (Jackson ImmunoResearch) were applied, followed by counterstaining with DAPI and mounting in antifade medium. Images were acquired using a Zeiss LSM 780 confocal microscope or Zeiss Axioscan.Z1 slide scanner.

### PTN-induced activation of human iPSC-derived microglia

Recombinant human PTN was expressed in *E. coli* Rosetta-gami 2 (DE3) cells and purified using sequential column chromatography on an ÄKTA system: a HiTrap SP FF cation-exchange column, a heparin column, and a Superdex 30 Increase 10/300 GL size-exclusion column. Finally, PTN-containing fractions were pooled and stored at −80 °C until use. SDS-PAGE analysis confirmed its high purity as a single predominant band.

The human induced pluripotent stem cells (hiPSC) were maintained on Matrigel (Corning)-coated plates in mTeSR™1 medium (STEMCELL Technologies) and passaged weekly using ReLeSR (STEMCELL Technologies). Induced microglia-like cells (iMG) were generated via a two-step differentiation protocol^69^, with minor modifications. Briefly, hiPSCs were differentiated into hematopoietic progenitor cells (HPCs) using the STEMdiff™ Hematopoietic Kit (STEMCELL Technologies). HPCs were then matured into microglia by culturing in a modified microglia differentiation medium containing DMEM/F-12 supplemented with B-27, N-2, Insulin-Transferrin-Selenium, non-essential amino acids, GlutaMAX, monothioglycerol, recombinant insulin, and the cytokines IL-34, TGF-β, and M-CSF for 24 d. For maturation, 100 ng/mL CD200 (PeproTech) and 50 ng/mL CX3CL1 (PeproTech) were added for 3 d. The mature iMG exhibited a characteristic microglial morphology and expressed microglial markers, as confirmed by immunocytochemistry for IBA1 (Abcam) and TMEM119 (Sigma-Aldrich). Then iMG were treated with PTN at different concentrations for stimulation. After treatment with PTN for 3 d, cells were harvested and lysed for proteomic profiling by LC-MS/MS.

Cells were lysed and digested as previously described ^33^. Approximately 50 ng of peptides were loaded onto a nanoscale C18 column (1.5 µm, 50 µm × 10 cm; HEB05001001518I-Rep) heated to 60 °C. Peptides were separated on a nanoElute 2 liquid chromatography system (Bruker), and MS data were acquired on a timsTOF Ultra2 mass spectrometer (Bruker) using dia-PASEF^70^. The PASEF acquisition range was set to 200-1700 m/z in MS1, with data-independent acquisition (DIA) MS2 scans acquired from 400-1200 m/z. Parameters included: cycle overlap = 4; mobilogram threshold = 50; MS/MS scans per cycle = 5; TOF resolution = 45,000; Exclusion window width was set to 0.15 m/z. Data were processed using the Pulsar directDIA workflow in Spectronaut (version 19.0). Carbamidomethylation (+57.02146 Da) was set as a fixed modification, and methionine oxidation (+15.99492 Da) as a variable modification, with up to two variable modifications allowed. The minimum peptide length was seven amino acids. Searches were conducted against the human UniProt FASTA database. Normalization and differential analysis were performed using the *limma* package. Proteins with an absolute z-score of log₂ fold change > 2, FDR < 0.05, and consistent directional changes across doses were considered differentially abundant.

## Resource availability

Raw data have been deposited in MassIVE under accession MSV000099715 (Username: MSV000099715_reviewer; Password: SCTP_PENG_2025).

## Code availability

The program of JUMP is publicly available from GitHub (https://github.com/JUMPSuite).

## Acknowledgements

We thank Dr. Takaomi Saido for providing the APP-KI mice. We thank the St. Jude Shared Resources and Core Facilities, including Animal Research Center, Flow Cytometry and Cell Sorting, Proteomics and Metabolomics. We also thank the insightful discussion with the Rossoll and Zhao groups at Mayo Clinic. This work was partially supported by National Institutes of Health grants RF1AG068581, R01AG092468, RF1AG064909, U19AG069701, and the ALSAC foundation.

## Author contributions

J.P., J.Y., Z.W., G.Y. conceived the project. X.Z., K.Y., D.L., L.H. performed FACS experiments. X.Z., K.Y. collected MS data. X.Z., H.S., Y.J. performed PL-based proteomics. Z.W., X.Z., X.R., Y.Z., Z.W. carried out PTN signaling experiments. P-C.C., Y.J., and K.H. supported IHC assays. Z.W., J.W., Z.W., A.A.H. supported MS analysis. X.Z., H.K.S., J.P., Z.F.Y., X.W., J.M.Y. performed computational analysis. J.Y. and Y.J. contributed to the scRNA-seq analysis. X.Z., H.K.S., and J.P. wrote the manuscript with input from all authors.

## Supplemental figures

**Figure S1.**
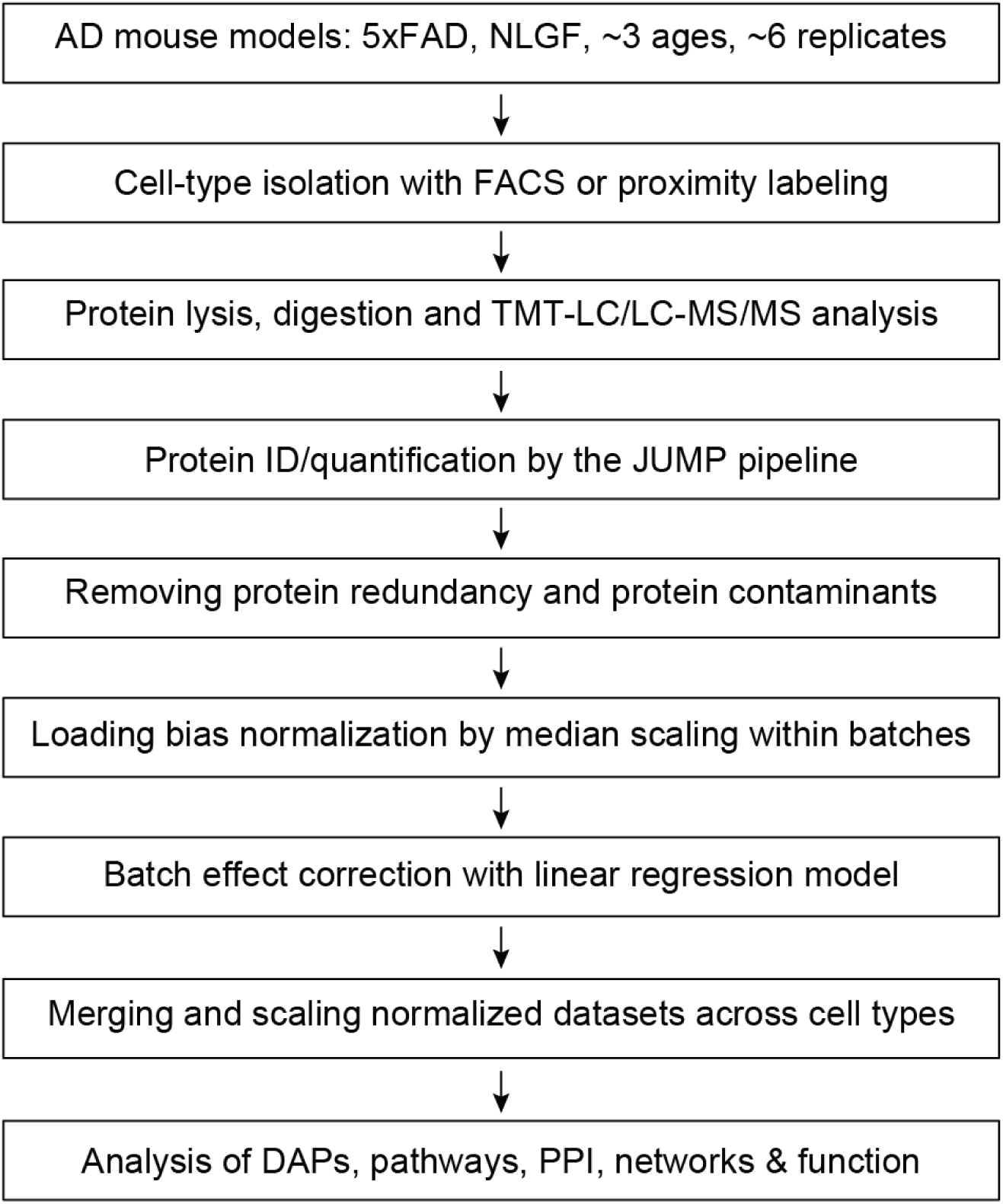
Cell type resolved proteomics data analysis workflow. Schematic of the experimental pipeline from AD mouse models through cell type isolation, TMT-based proteomics, and data processing.

**Figure S2.**
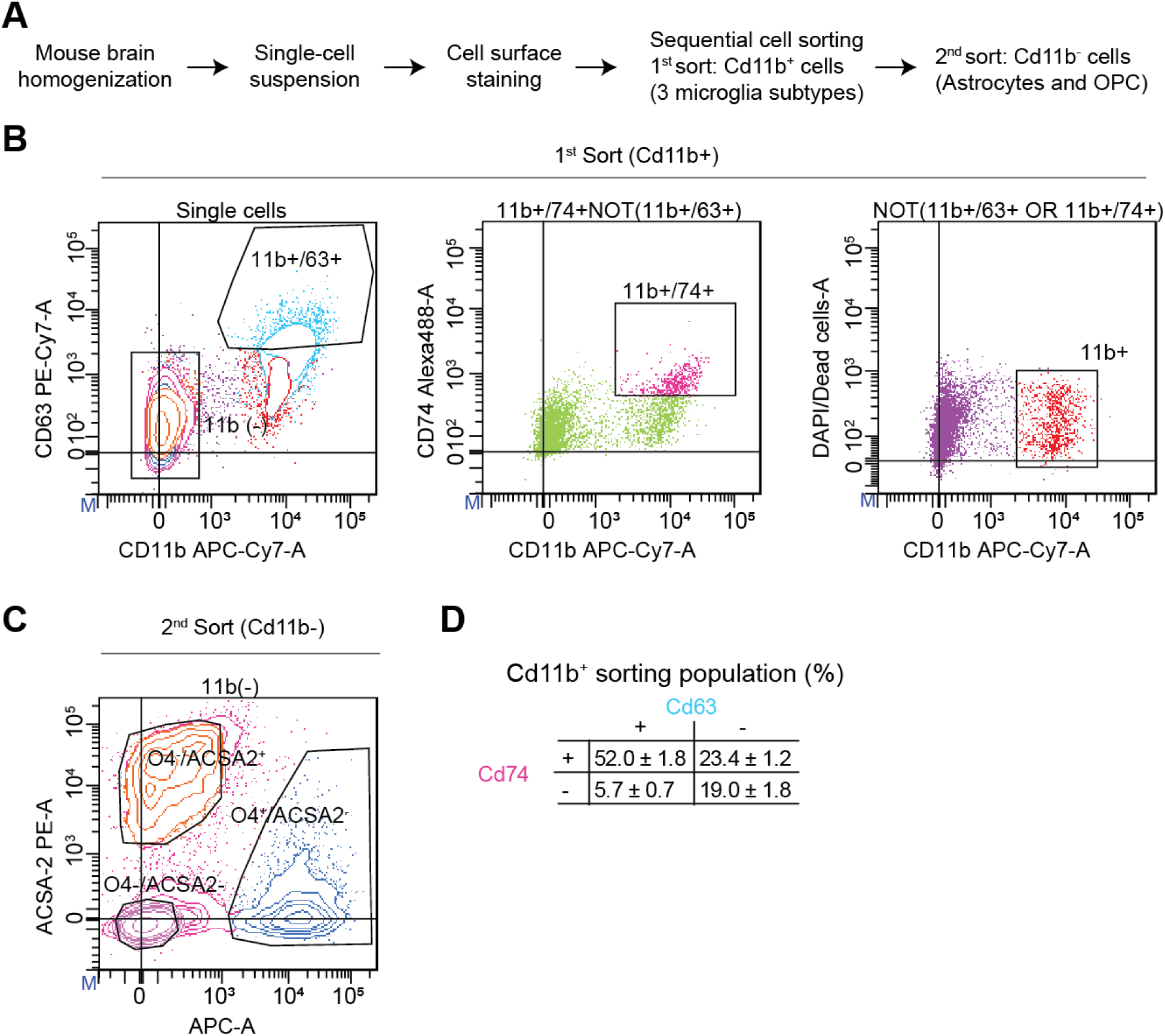
FACS-based strategy to sort non-neuronal brain cell types. **(A)** Workflow for isolating non-neuronal cells from mouse brain. **(B-C)** Sequential FACS gating strategy to sort Cd11b⁺ microglia subtypes (MG1-MG3) and Cd11b⁻ population: astrocytes and OPC. **(D)** Fractional percentages of Cd11b⁺ sorted cells obtained from different gating strategies. Data are presented as mean ± standard error of the mean (SEM) from 8-month-old FAD mice (*n* = 6). As the population of Cd11b⁺/Cd63⁺/Cd74⁻ was small, we decided to merge Cd11b⁺/Cd63⁺/Cd74⁻ and Cd11b⁺/Cd63⁺/Cd74⁺ into one subtype of MG3 (Cd11b⁺/Cd63⁺).

**Figure S3.**
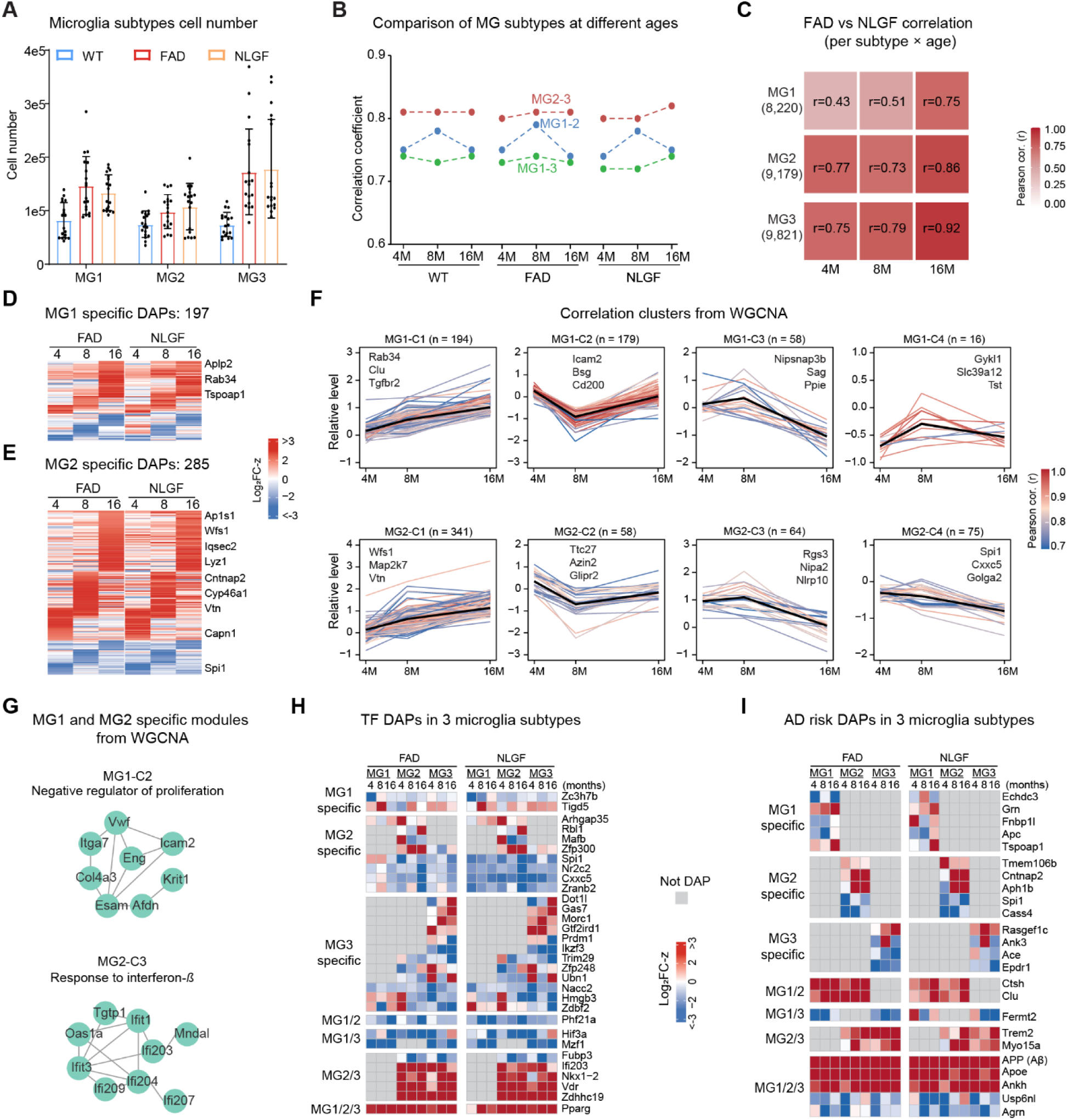
Microglia subtype proteomic characterization in AD mice. **(A)** Quantification of MG1, MG2, and MG3 cell numbers in WT, 5xFAD, and NLGF mice (*n* = 6 per group). **(B)** Pairwise correlations of three MG subtypes at each age group in both AD mice, using z-transformed MS intensities. **(C)** FAD-NLGF proteomic correlation of each subtype at different ages. **(D)** MG1-specfic DAPs across ages in two AD mice showing consistent patterns. **(E)** MG2-specfic DAPs. **(F)** Protein clusters in MG1 or MG2 DAPs identified by WGCNA. Each plot shows the relative abundance of proteins within a cluster; individual lines represent single proteins, and the black line indicates the cluster eigengene trend. **(G**) Representative PPI modules enriched in MG1 and MG2 subtypes. **(H,I)** Proteomic changes of TFs and AD risk genes that are consistent in both AD mice, including those specific to MG1, MG2, MG3, and shared among subtypes.

**Figure S4.**
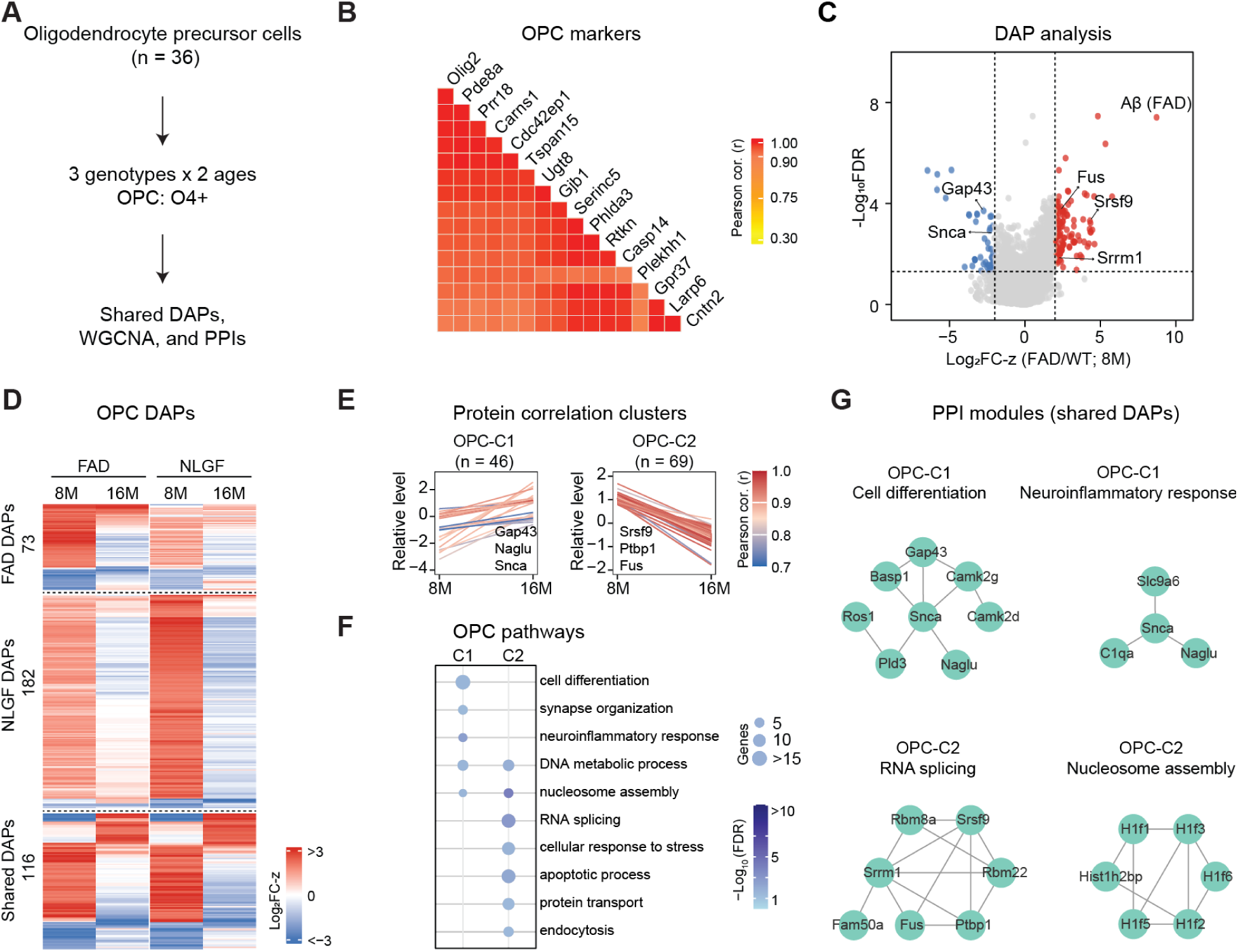
Proteomic characterization of OPC in AD models. **(A)** Workflow. Two ages were analyzed, as early-stage samples failed quality control due to suboptimal instrument performance. **(B)** Pearson correlation heatmap of top OPC-specific marker proteins, ranked by their correlation with Olig2, the target of the O4⁺ antibody. **(C)** Representative volcano plot from OPC at 8 months comparing FAD with WT. **(D)** Heatmaps of age-dependent DAPs grouped by genotypes: FAD-, APPKI-specific, and shared by both models**. (E)** Two OPC protein clusters identified through WGCNA. Each line represents a single protein’s trajectory over time. **(F)** Pathway enrichment of the two OPC protein clusters. Dot size corresponds to the number of overlapping genes, and color intensity reflects the statistical significance. **(G)** Representative PPI modules enriched in OPC proteins clusters.

**Figure S5.**
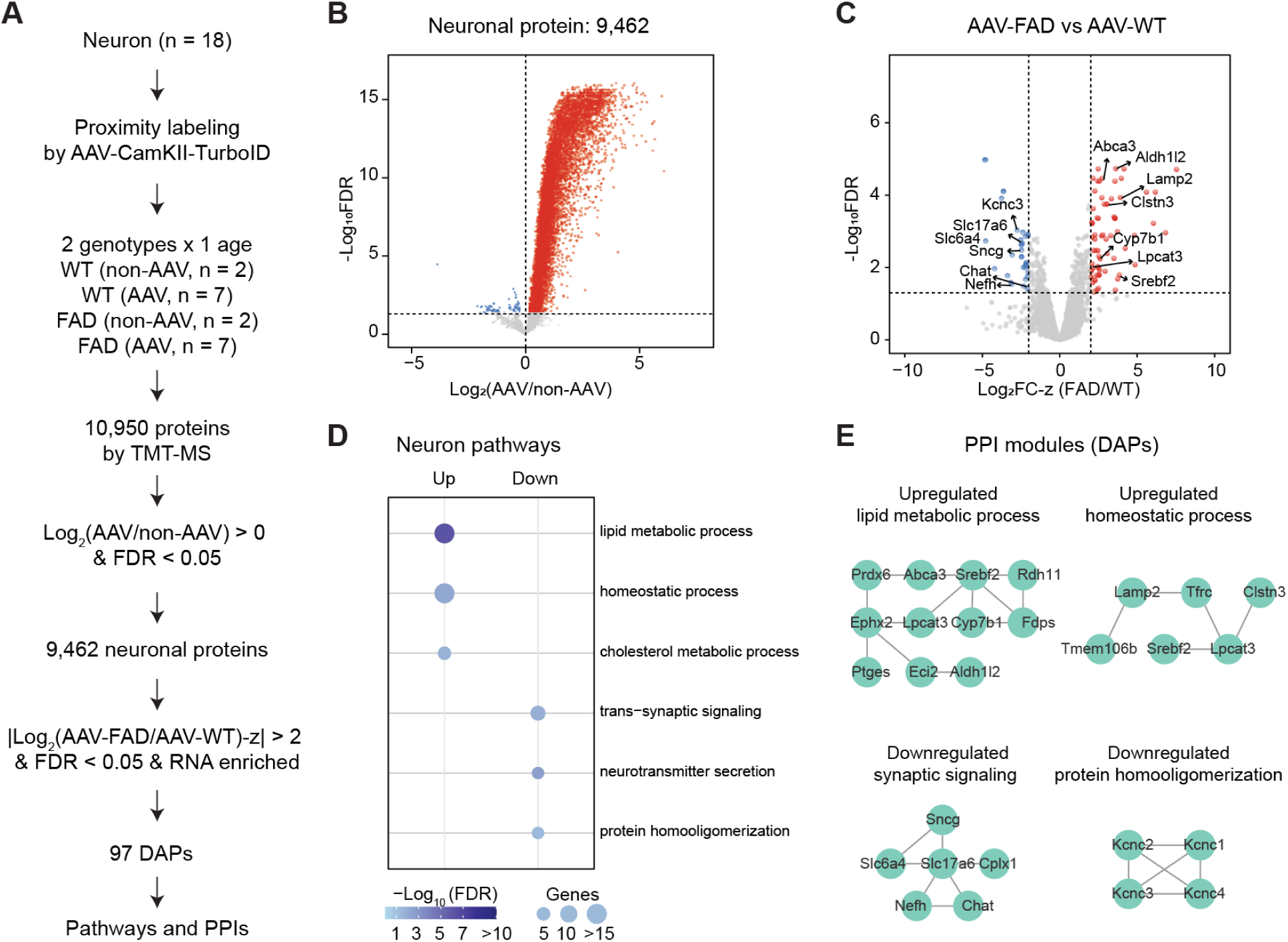
Deep neuron-specific proteomics reveals AD-associated alterations. **(A)** Workflow for neuronal proteomics using AAV-mediated TurboID-CaMKII proximity labeling in neurons. A total of 10,950 proteins were quantified, with 9,462 identified as neuron-enriched and 97 DAPs subjected to further analysis. **(B)** Volcano plot of AAV-enriched proteins compared to non-AAV background proteome. **(C)** Volcano plot of DAPs between FAD and WT in AAV enriched proteins. **(D)** Pathway enrichment of upregulated and downregulated DAP. Dot size corresponds to the number of overlapping genes, and color intensity reflects the statistical significance**. (E)** Representative PPI modules enriched in upregulated and downregulated neuronal DAPs.

**Figure S6.**
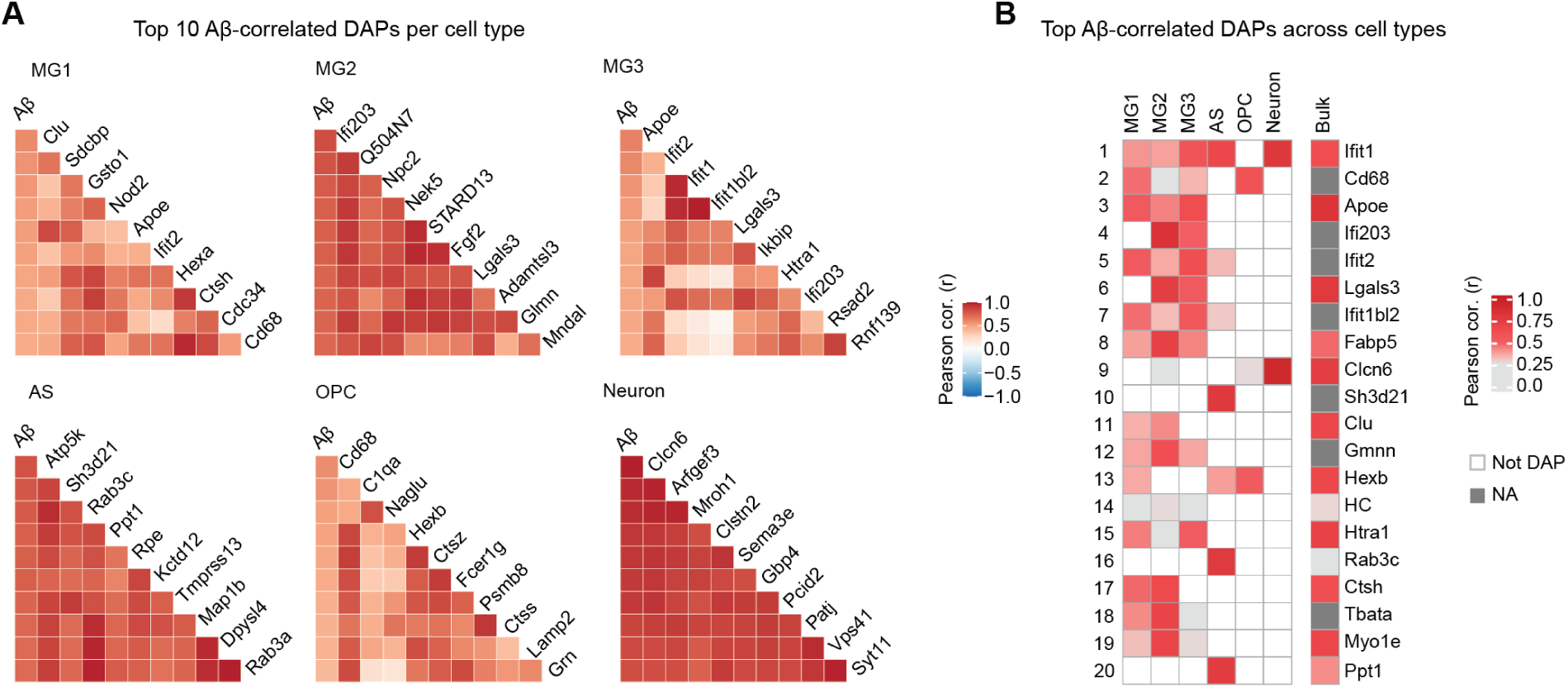
Aβ-correlated protein signatures across brain cell types in FAD. **(A)** Heatmaps showing the top 10 cell type–specific DAPs correlated with the Aβ peptide LVFFA***E***DVGSNK. **(B)** Heatmap of the top 20 Aβ-correlated DAPs across cell types, ranked by an order summed from Aβ-correlation values in individual cell types. The correlation of these DAPs in bulk proteomics was also shown.

**Figure S7.**
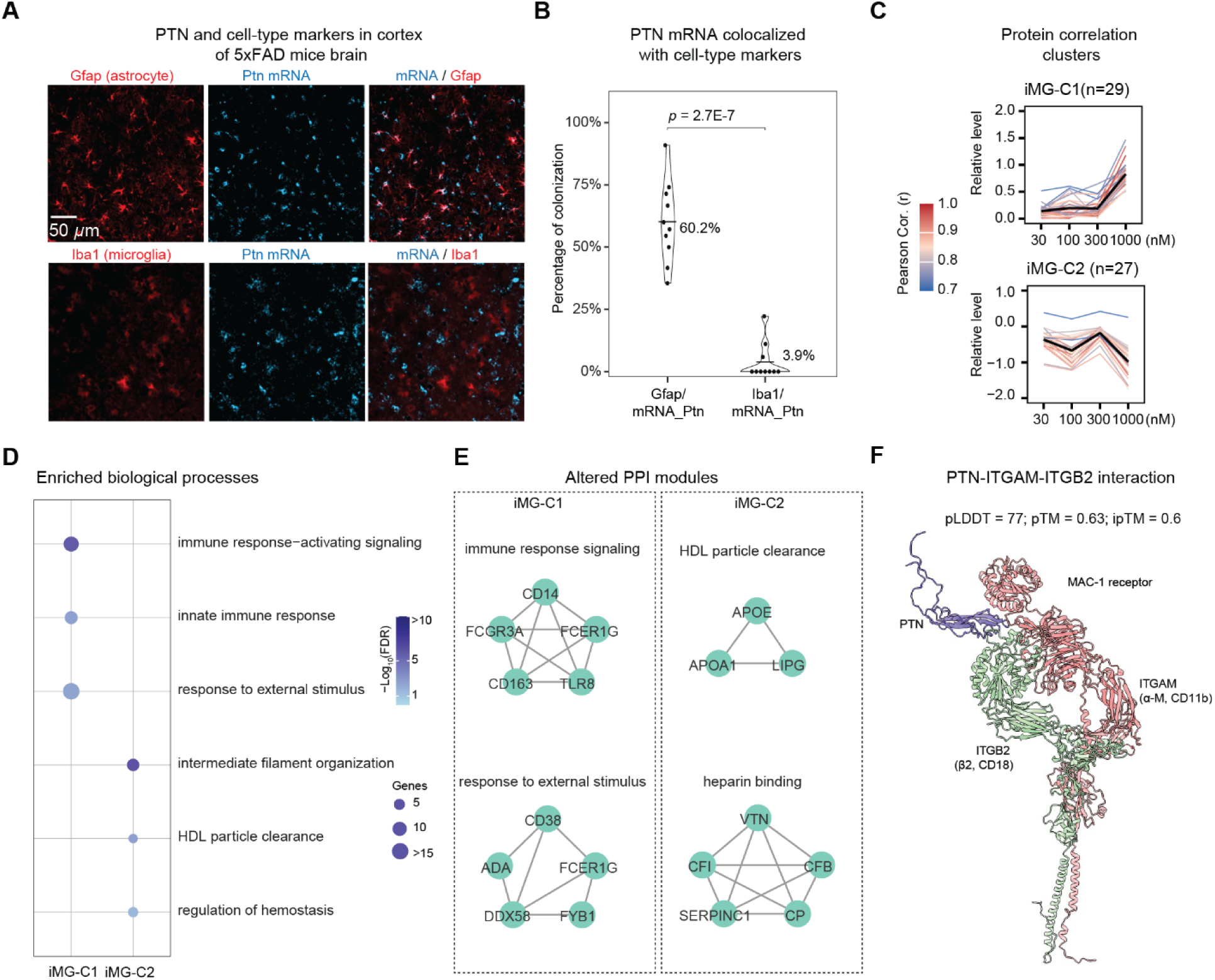
PTN-responsive protein programs and a putative MAC-1 interaction. **(A**) In situ localization of *Ptn* and cell type markers in 5xFAD cortex. RNAscope/IF for Gfap (astrocytes, red) or Iba1 (microglia, red) with **Ptn mRNA** (cyan). Merged images at right. Scale bar, 50 µm. **(B)** Quantification of Ptn mRNA colocalization with cell type markers showing predominant enrichment in astrocytes (Gfap, 60.2%) and minimal overlap with microglia (Iba1, 3.9%) (*p* = 2.7 × 10⁻⁷). **(C)** Two PTN-induced protein clusters identified through WGCNA. Each line represents a single protein’s trajectory over time. **(D)** Pathway enrichment of the two PTN-induced microglia protein clusters. Dot size corresponds to the number of overlapping genes, and color intensity reflects the statistical significance. **(E)** Representative PPI modules altered within WGCNA modules. **(F)** AlphaFold-Multimer model suggesting a PTN-MAC-1 (ITGAM/ITGB2) interface. Reported confidence metrics: pLDDT = 77, pTM = 0.63, ipTM = 0.60 (illustrative only).

## Supplemental tables

**<u>Table S1</u>**

Table S1.1. Summary of sample information for different cell types.

Table S1.2. Detailed sample information utilized for deep in vivo cell-type-resolved proteomic profiling of the AD mouse brain.

Table S1.3. Proteomic profiling of mouse brain cell types by TMT-LC/LC-MS/MS Table S1.4. List of cell-type markers utilized as positive controls.

Table S1.5. Comprehensive list of differentially abundant proteins identified across cell types and conditions

**<u>Table S2</u>**

Table S2.1. Comparison of mouse AD and WT proteomes in MG1 at multiple ages

Table S2.2. Comparison of mouse AD and WT proteomes in MG2 at multiple ages

Table S2.3. Comparison of mouse AD and WT proteomes in MG3 at multiple ages

Table S2.4. Mouse DAPs of 3 MG subtypes with TF and risk genes highlighted

Table S2.5. Top microglia subtype specific markers in mice

Table S2.6. Pathway enrichment of DAPs in 3 MG subtypes in mice

Table S2.7. WGCNA modules of MG1 in mice

Table S2.8. Pathway enrichment analysis of distinct WGCNA modules in MG1

Table S2.9. WGCNA modules of MG2 in mice

Table S2.10. Pathway enrichment analysis of distinct WGCNA modules in MG2

Table S2.11. WGCNA modules of MG3 in mice

Table S2.12. Pathway enrichment analysis of distinct WGCNA modules in MG3

**<u>Table S3</u>**

Table S3.1. Differential analysis of astrocytes across three ages

Table S3.2. Astrocyte DAP change patterns across ages and genotypes

Table S3.3. Top astrocytes specific markers in mice

Table S3.4. Pathway enrichment of genotype shared DAPs in astrocytes in mice

Table S3.5. WGCNA modules of astrocytes in mice

Table S3.6. Pathway enrichment analysis of distinct WGCNA modules in astrocytes

**<u>Table S4</u>**

Table S4.1. Differential analysis of oligodendrocyte precursor cell (OPC) across two ages

Table S4.2. OPC DAP change patterns across ages and genotypes

Table S4.3. Top OPC specific markers in mice

Table S4.4. Pathway enrichment of genotype shared DAPs in OPC in mice

Table S4.5. WGCNA modules of oligodendrocyte precursor cell

Table S4.6. Pathway enrichment analysis of distinct WGCNA modules in OPC

**<u>Table S5</u>**

Table S5.1. Differential analysis of neuron Table S5.2. Neuronal DAPs

Table S5.3. Pathway enrichment of up- and down-regulated neuronal DAPs

**<u>Table S6</u>**

Table S6.1. MEGENA module summary based on the combined proteins from all cell types

Table S6.2. MEGENA module summary of MG1

Table S6.3. MEGENA module summary of MG2

Table S6.4. MEGENA module summary of MG3

Table S6.5. MEGENA module summary of astrocyte (AS)

Table S6.6. MEGENA module summary of oligodendrocyte precursor cell (OPC)

Table S6.7. MEGENA module summary of Neuron

Table S6.8. MEGENA hub comparison across the cell types

**<u>Table S7</u>**

Table S7.1. Pathway enrichment analysis of mouse whole brain proteome–specific proteins absent from cell-type proteomes

Table S7.2. Comparison of cell-type proteomes with mouse brain bulk proteome

Table S7.3. Integrative ranking of genes/proteins in microglia

Table S7.4. Integrative ranking of genes/proteins in astrocyte

Table S7.5. Integrative ranking of genes/proteins in OPC

Table S7.6. Integrative ranking of genes/proteins in Neuron

Table S7.7. Integrative ranking of top genes/proteins in each cell type

Table S7.8. Integrated rank of Aβ correlations from cell-type DAPs

**<u>Table S8</u>**

Table S8.1. Cell-type expression from single-cell transcriptomes across three ages of FAD mice

Table S8.2. Cell-type proteome compared with transcriptome

Table S8.3. DAP consistency UniProt location and RNA cell type enrichment

Table S8.4. Intercellular signaling proteins derived from secreted DAPs

**<u>Tables S9</u>**

Table S9.1. Proteomic profiling of PTN-treated human iPSC-derived microglia at different doses by DIA-LC–MS/MS

Table S9.2. Consistently regulated DAPs across PTN doses in human iPSC-derived microglia

Table S9.3. GO BP enrichment of consistently regulated DAPs across PTN doses

Table S9.4. WGCNA analysis of PTN dose-dependent treatment in iPSC-derived microglia

Table S9.5. GO enrichment analysis of WGCNA modules from PTN dose-dependent treatment in iPSC-derived microglia

Table S9.6. Consistent protein changes between human iPSC-derived microglia and MG3 DAPs in AD mice

